# Interaction of *G2019S LRRK2* and metabolic syndrome in a two-hit mouse model of Parkinson’s disease: LRRK2-driven systemic depletion of pyrimidine nucleosides

**DOI:** 10.64898/2025.12.31.697225

**Authors:** Yue Ma, Madalynn L. Erb, Kayla A. Sipple, Alina V. Offerman, Junwei Niu, Zachary B Madaj, Zhen Fu, Kin H Lau, Darren J. Moore

## Abstract

*G2019S LRRK2* is the most common cause of familial Parkinson’s disease (PD) and is associated with sporadic PD, arising from the interplay of genetic predisposition, environmental exposure and aging. Metabolic syndrome is implicated as a risk factor for PD, but the interaction between *G2019S LRRK2* and metabolic stress in disease pathogenesis remains unclear. We employed high-fat diet (HFD) feeding to induce metabolic syndrome in aged mutant *LRRK2* mice, followed by system-wide characterization, including metabolomic or proteomic profiling, and bulk or single-nucleus RNA sequencing. We find that thymidine and deoxyuridine levels are consistently reduced across tissues in *G2019S LRRK2* knockin mice accompanied by increased hepatic expression of thymidine phosphorylase. HFD exposure further unmasks disruptions in purine and energy metabolism in brain and lung of *G2019S LRRK2* knockin mice, with midbrain astrocytes and oligodendrocytes exhibiting the most pronounced impairment in oxidative phosphorylation transcriptional pathways. Our findings demonstrate that pre-existing metabolic syndrome unmasks widespread disruptions in systemic nucleotide and energy metabolism and exacerbates mitochondrial dysfunction in *G2019S LRRK2* knockin mice. This conditional “two-hit” phenotype underscores the critical role of environmental factors, such as diet, in revealing metabolic vulnerabilities associated with PD-linked genetic backgrounds, and provides potential metabolic targets for therapeutic intervention in PD.

## Introduction

Parkinson’s disease (PD) is the most common neurodegenerative movement disorder, affecting 1-2% of people over 65 years old, characterized by the progressive loss of dopaminergic neurons in the substantia nigra pars compacta and by the accumulation of intracellular proteinaceous inclusions called Lewy bodies [1]. The loss of dopaminergic innervation results in a range of clinical features including tremor, bradykinesia, rigidity and postural instability [1]. Among all PD cases, about 10% are familial, which are linked to mutations in >20 distinct genes [1, 2]. Mutations in the *leucine-rich repeat kinase 2* (*LRRK2*) gene are the most common cause of late-onset, autosomal dominant familial PD (*LRRK2*-PD), particularly the *G2019S LRRK2* mutation, which is the most common and accounts for 4% of familial and 1% of sporadic PD cases worldwide [3–5]. However, the penetrance of the G2019S mutation is incomplete, suggesting that additional modifiers, such as diet or metabolic stress, may be required to unmask disease-related phenotypes.

Metabolic syndrome (MetS) is a cluster of conditions that often occur together, including obesity, hyperglycemia, hypertension, and dyslipidemia [6], and having just one of these components increases the risk of heart disease, stroke, and type-2 diabetes [6]. MetS affects one-third of U.S. adults, and the prevalence of MetS increases rapidly with age [7]. A nationwide cohort study found that MetS and its components are independent risk factors for the development of PD [8]. Another cohort-based comparative clinical analysis has shown prediabetes is more frequent among individuals with *LRRK2* mutations, suggesting a connection between LRRK2 and metabolic dysfunction [9]. However, it is currently unclear whether pre-existing MetS enhances neurotoxic pathways in *LRRK2*-PD, or whether mutant LRRK2 contributes to the development of MetS. Understanding this relationship will provide critical insights into disease pathways, allowing us to evaluate whether targeting metabolic pathways in PD subjects represents a promising strategy for modulating neuropathology and neurodegeneration.

Here, we use a high-fat diet (HFD) to induce metabolic syndrome in a mutant LRRK2 mouse model of PD and systematically evaluate whether metabolic stress unmasks latent vulnerabilities that contribute to PD pathogenesis. Moreover, the application of multi-omic strategies offers opportunities to identify candidate biomarkers of *LRRK2*-PD progression and to reveal potential therapeutic entry points for genetically at-risk individuals.

## Results

### Prolonged high-fat diet exposure results in metabolic syndrome features in *G2019S LRRK2* knockin and wild-type mice

Modifying diet composition can induce multiple features of metabolic syndrome (MetS) in rodent models. However, the effects of specific diets on individual metabolic pathways varies with genetic background, causing different strains of mice to respond differently to specific diets [10]. To examine the effect of high-fat diet (HFD) on *G2019S LRRK2* knockin mice, 11 month-old homozygous *G2019S LRRK2* knockin (KI/KI) mice and their wild-type (WT) littermates were fed a HFD (60% of calories from fat) or control diet (Con. diet, 10% of calories from fat) for 5 months (**Fig. 1a**). As expected, we find that bodyweight is markedly increased in HFD-fed mice compared to those on control diet, although *G2019S LRRK2* mice gain modestly less weight than WT mice (**Fig. 1b**). We also observe elevated blood glucose levels (**Fig. 1c**) and increased plasma insulin levels (**Fig. 1d**) in HFD-fed mice. To examine the impact of a high-fat diet on lipid levels, we performed lipidomic analysis across multiple tissues. We find that plasma lipid levels are significantly elevated in HFD-fed mice (**Fig. 1e, f**, **Table S1**). Additionally, lipid composition is altered across multiple tissues (liver, brain, lung, kidney) in HFD-fed mice (**Fig. S1**, **Table S1**). However, we do not observe significant differences in lipid composition between *G2019S LRRK2* knockin mice and their WT littermates under HFD or control diet across tissues (**Fig. S2**, **Table S1**). To investigate the effects of a high-fat diet on hepatic steatosis, we analyzed liver tissue using H&E staining to assess lipid accumulation. We find evidence of both macrovesicular and microvesicular steatosis in the livers of HFD-fed mice (**Fig. 1g**). Quantitation reveals that the percentage of lipid-positive area in HFD-fed mouse liver is significantly higher than that in control diet-fed mice (**Fig. 1h**). These data demonstrate that HFD exposure equivalently induces key biochemical and metabolic features of MetS in WT and *G2019S LRRK2* KI mice over 5 months.

**Figure 1.**
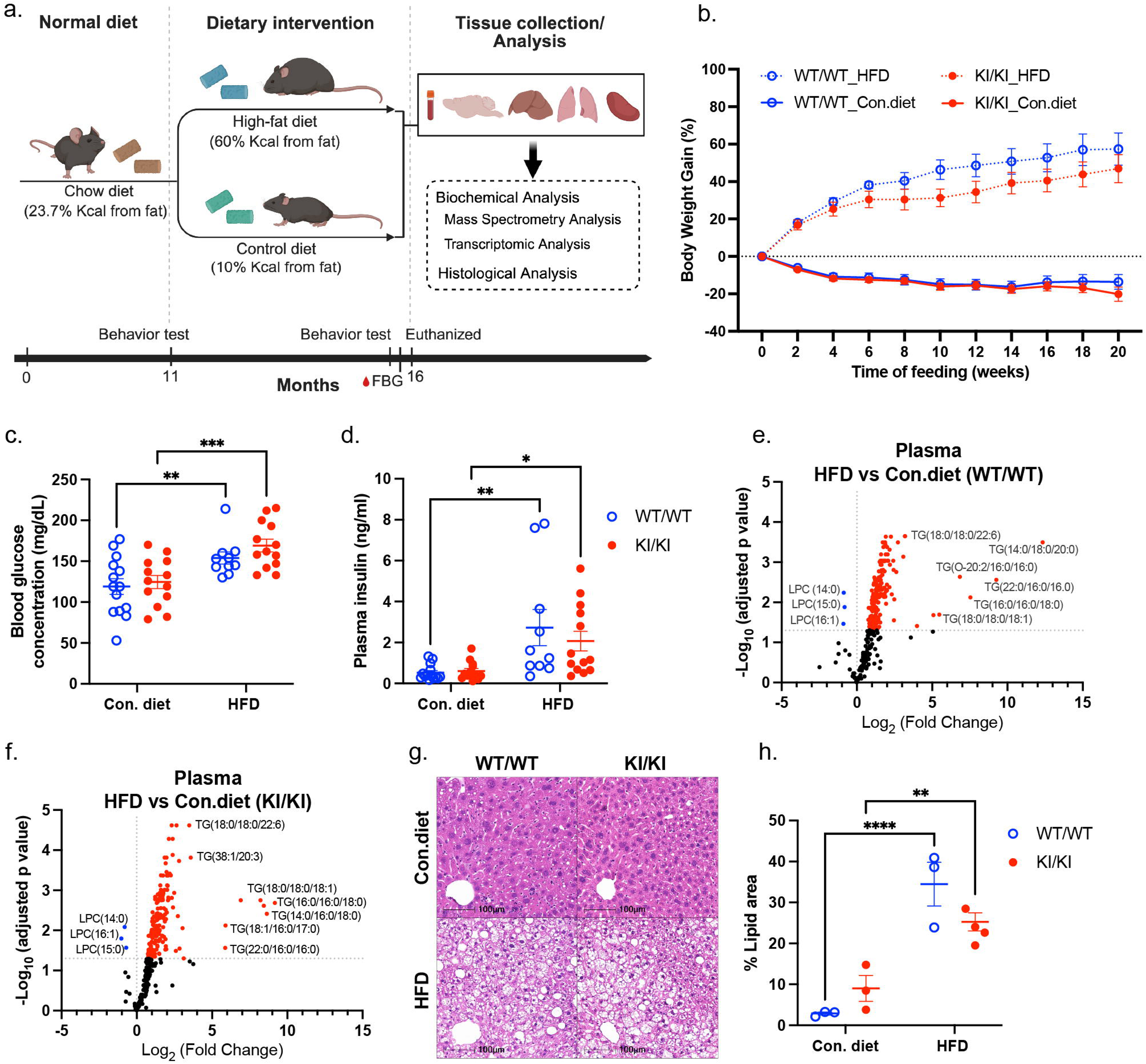
Characterization of diet-induced metabolic syndrome in *G2019S LRRK2* knockin mice. **a** Schematic diagram of the experimental design. Mice were fed with normal chow diet (23.7% Kcal from fat) for 11 months, then they were randomly divided into high-fat diet (60% Kcal from fat) or control diet (10% Kcal from fat) groups, the mice were kept on their specific diets for an additional five months. Behavioral tests were conducted before the diet switch and again four months afterward. Body weight was measured biweekly. Fasting blood glucose and plasma insulin levels were measured two weeks prior to euthanasia. At the age of 16 months old, plasma, brain, liver, lung, and kidney were collected and subjected to biochemical and histological analysis. **b** Body weight changes along with 20 weeks of specific diet feeding (*n* = 10-14 per group). **c** Blood glucose and **d** Plasma insulin level at 20 weeks post specific diet feeding (*n* = 10-14 per group). **e-f** Volcano plot showing differences in the abundance of lipids between HFD and Con. diet in the plasma of *G2019S LRRK2* knockin mice and wild-type controls, each point represents a lipid species. X-axis shows log_2_ (fold change) in lipid abundance between high-fat diet and control diet groups. Y-axis shows -log_10_ (adjusted *p*-value) from limma’s empirical Bayes moderated analysis (eBayes) combined with Benjamini-Hochberg (BH) for multiple testing correction (*n* = 6 per group). Lipids with adjusted *P*<0.05 are considered significantly altered. Significantly upregulated lipids are highlighted in red and significantly downregulated lipids are highlighted in blue. **g** Representative images of Hematoxylin and Eosin (H&E) staining of the livers collected at the terminal stage, scale bar: 100 µm. **h** Quantification of lipid area in H&E-stained liver sections (*n* = 3-4 per group). WT/WT, wild-type mice; KI/KI, homozygous *G2019S LRRK2* knockin mice; TG, triglycerides; LPC, lysophosphatidylcholine. Data are presented as mean ± SEM. A two-way ANOVA was performed to evaluate the effects of high-fat diet and *G2019S LRRK2* mutant and their interaction followed by Fisher’s LSD comparisons. Significant main effects were found (\**P*<0.05, \*\**P*<0.01, \*\*\**P*<0.001, \*\*\*\**P*<0.0001).

### Pre-existing metabolic syndrome induces anxiety-like behavior in *G2019S LRRK2* knockin and wild-type mice

*G2019S LRRK2* knockin mice are reported to exhibit abnormalities in striatal function and increased hyperphosphorylated tau (pSer202/Thr205) at 12 months of age [11]. Surprisingly, these mice do not experience motor impairment or neurodegeneration in the substantia nigra pars compacta (SNpc) at this age [11]. To test the effect of pre-existing MetS on motor deficits, we conducted behavioral assessments using the open field test (Fig. 2a) and CatWalk digital gait analysis. We find that the total distance traveled in open field is significantly decreased in HFD-fed *G2019S LRRK2* knockin mice compared to control diet, and HFD-fed WT mice show a similar non-significant decrease (**Fig. 2b**). We do not observe any differences among groups in the distance traveled in the center zone relative to the total distance or in the number of entries (**Fig. 2c-d**). CatWalk gait analysis reveals a significant reduction in average speed in HFD-fed *G2019S LRRK2* knockin mice relative to those fed a control diet, with a similar non-significant reduction in HFD-fed WT mice (**Fig. 2e**). We also observe that stand time was significantly increased in HFD-fed mice compared to those fed a control diet (**Fig. 2f-i**), while no differences are detected in cadence or base of support (**Fig. 2j-l**). To assess anxiety-like behavior in these mice, we conducted the elevated plus maze test (**Fig. 2m**). In this test, HFD-fed mice spend significantly less time in the open arms compared to control diet-fed mice (**Fig. 2n**). Additionally, the number of entries into the open arms and the ratio of distance travelled in open arms and center zone relative to total distance are reduced in HFD-fed mice compared to controls (**Fig. 2o-p**), indicating increased anxiety-like behavior. To test the effect of pre-existing MetS on cognitive deficits, we conducted Y-maze tests to assess spatial working and reference memory, although no significant differences in alternation behavior are observed among groups (**Fig. 2q-r**). These data indicate that HFD exposure results in reduced locomotor activity and increase anxiety-like behavior in mice, but without obvious differences between *LRRK2* genotypes.

**Figure 2.**
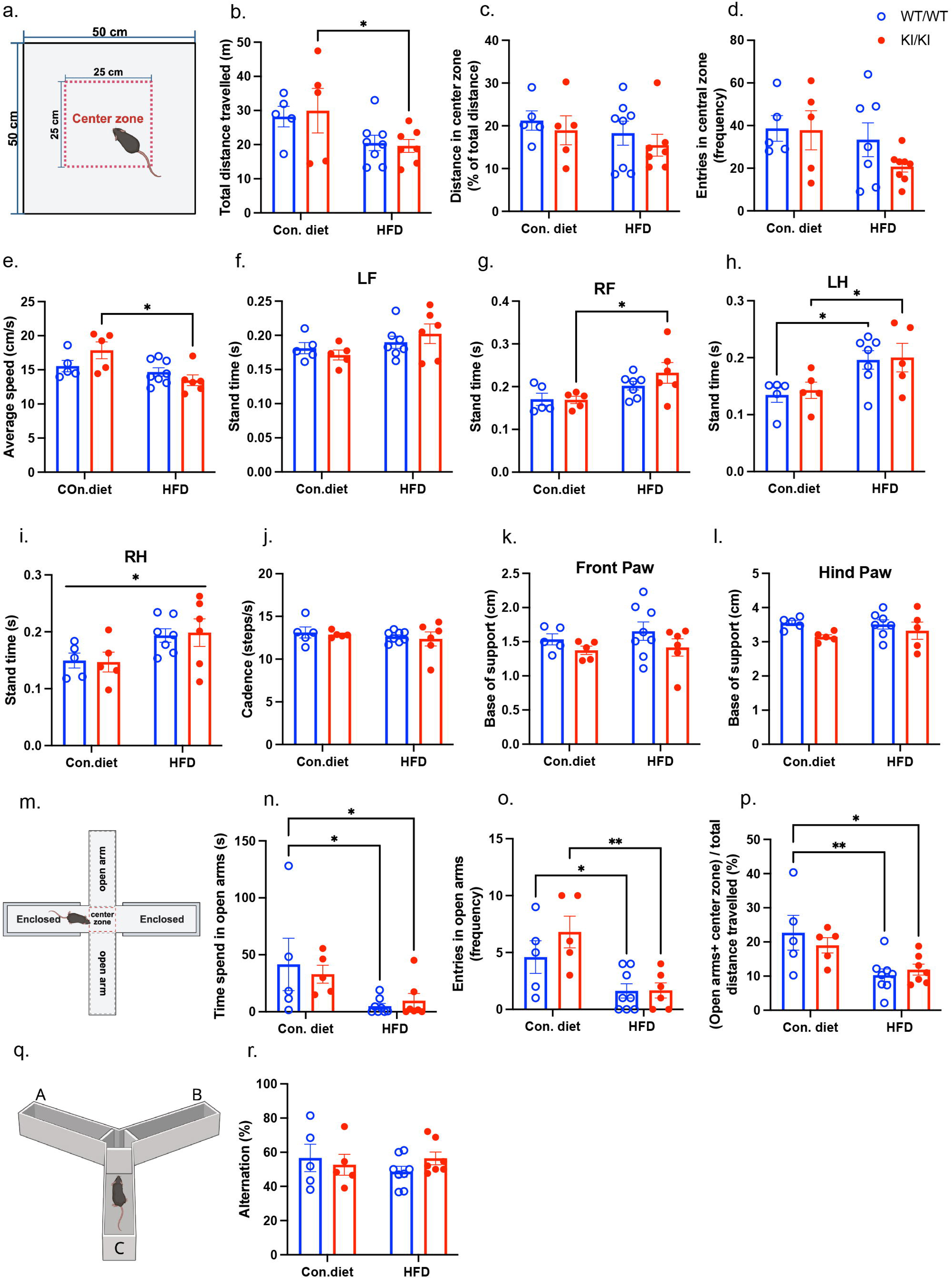
Behavior deficits associated with pre-existing metabolic syndrome in *G2019S LRRK2* knockin mice. Behavioral tests were conducted four months after the initiation of diet switch (*n* = 5-8 per group). **a-d** Open field test. **a** Scheme and dimensions of the open field apparatus; **b** Total distance travelled (m) during the 15-minute open field test; **c** Distance travelled in center zone of total distance of the open field test during 15-minute session; **d** Entries in central zone (frequency) during the 15-minute open field test. **e-l** Catwalk gait analysis. **e** Average speed (cm/s); **f-i** Stand time of each paw (s), LF, left front; RF, right front; LH, left hind; RF, right hind. **j** Cadence (steps/s); **k, i** Base of support; **m-p** Elevated plus maze. **m** Scheme and annotation of the elevated plus maze apparatus; **n** Time spend (s) in open arm of elevated plus maze during the 10-minute test; **o** Entries in open arms (frequency) of elevated plus maze during the 10-minute test; **p** Ratio of distance traveled in the open arms and center zone relative to the total distance during the elevated plus maze. **q-r** Y-maze. **q** Scheme and annotation of the Y-maze apparatus; **r** Spontaneous alternation (%) calculated in the 10-minute Y-maze test. WT/WT, wild-type mice; KI/KI, homozygous *G2019S LRRK2* knockin mice. Data are presented as mean ± SEM. A two-way ANOVA was performed to evaluate the effects of high-fat diet and *G2019S LRRK2* mutant and their interaction followed by Fisher’s LSD comparisons. Significant main effects were found (\**P*<0.05, \*\**P*<0.01).

### Pre-existing metabolic syndrome induces gliosis in *G2019S LRRK2* knockin and wild-type mice

A high-fat diet can induce inflammation in the hypothalamus, a brain region critical for regulating energy balance [12]. Notably, hypothalamic inflammation occurs before significant body weight gain and involves the rapid activation of a complex network of cells [12, 13]. This neuroinflammation is closely linked to the development of obesity and related metabolic disorders [13]. To evaluate the effect of pre-existing MetS on neuroinflammation across the brain in *G2019S LRRK2* knockin mice, we assessed reactive gliosis by immunostaining for the microglial marker Iba1 (**Fig. 3a, d, g**) or the astrocyte marker GFAP (**Fig. 3j**, **Fig. S3a, c**). We observe an increased number of Iba1-positive microglia in the striatum (**Fig. 3b**) and substantia nigra (**Fig. 3e**) of HFD-fed mice. However, an increased soma size of microglia is only observed in the striatum of HFD-fed WT mice (**Fig. 3c, f**), which indicates microglial activation. Additionally, both the number and soma size of microglia are elevated in the hypothalamus of HFD-fed WT mice but not G2019S *LRRK2* mice (**Fig. 3h-j**). We also observe an increased percentage of GFAP-positive astrocytes in the hypothalamus of HFD-fed *G2019S LRRK2* knockin mice compared to those on a control diet and a similar non-significant increase in HFD-fed WT mice (**Fig. 3k**). However, we do not observe any difference in the percentage of GFAP-positive astrocytes in HFD relative to control diet in the striatum or substantia nigra (**Fig. S3a-d**). Our data indicate that HFD exposure induces reactive gliosis in the mouse brain, particularly the hypothalamus, with *G2019S LRRK2* KI mice displaying a modest reduction in HFD-induced microglial activation relative to WT mice. These findings might suggest that G2019S LRRK2 modestly compromises the microglial activation response.

**Figure 3.**
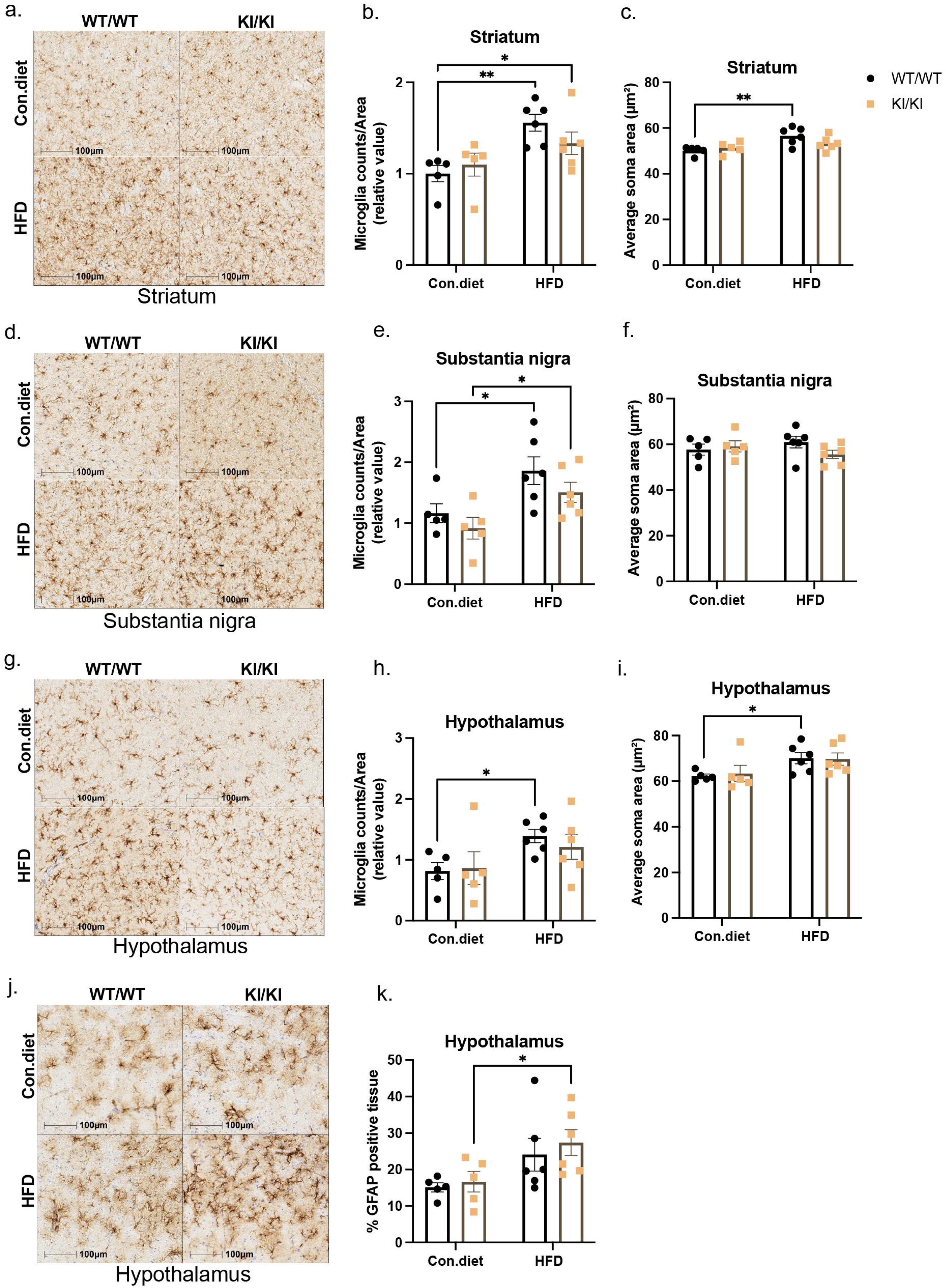
Pre-existing metabolic syndrome induces reactive gliosis in *G2019S LRRK2* knockin mice. **a, d, g** Representative images of Iba1 immunostaining in the striatum (**a**), substantia nigra (**d**), and hypothalamus (**g**) showing microglial distribution. Scale bar: 100 µm. **b, e, h** Quantification of Iba1-positive microglial counts relative to area in the striatum (**b**), substantia nigra (**e**), and hypothalamus (**h**). **c, f, i** Quantification of average microglial soma area (µm²) in the striatum (**c**), substantia nigra (**f**), and hypothalamus (**i**) (*n* = 5-6 per group). **j** Representative images of GFAP immunostaining in the hypothalamus showing astrocytes distribution. Scale bar: 100 µm. **k** Quantification of GFAP-positive area in the hypothalamus (*n* = 5-6 per group). WT/WT, wild-type mice; KI/KI, homozygous *G2019S LRRK2* knockin mice. Data are presented as mean ± SEM. A two-way ANOVA was performed to evaluate the effects of high-fat diet and *G2019S LRRK2* mutant and their interaction followed by Fisher’s LSD comparisons. Significant main effects were found (\**P*<0.05, \*\**P*<0.01).

### Pre-existing metabolic syndrome unmasks striatal dopamine loss without nigrostriatal dopaminergic neuronal loss in *G2019S LRRK2* knockin mice

Dopaminergic neurons in the substantia nigra pars compacta (SNpc) express tyrosine hydroxylase (TH) and project their axons to the striatum [14]. To assess the effect of pre-existing MetS on dopaminergic neurodegeneration in the SNpc, we performed unbiased stereological cell counting of TH-positive dopaminergic neurons and total Nissl-positive neurons. However, we do not detect nigral neuronal loss caused by G2019S LRRK2, HFD or their combination (**Fig. 4a-c**). To evaluate TH-positive innervation of the striatum, the percentage of TH-positive tissue was measured. Intriguingly, we find a significant decrease of TH-positive nerve terminals in HFD-fed *G2019S LRRK2* knockin mice compared to WT mice under control diet (**Fig. 4d-e**), and there is also a similar non-significant reduction in *G2019S LRRK2* KI mice fed a control diet. Additionally, we performed targeted metabolic profiling of striatal tissue by LC-MS and find a significant decrease of dopamine (DA) levels selectively in HFD-fed *G2019S LRRK2* knockin mice compared to other groups (**Fig. 4f**), indicating an interaction of G2019S LRRK2 and HFD on the density and function of striatal dopaminergic nerve terminals.

**Figure 4.**
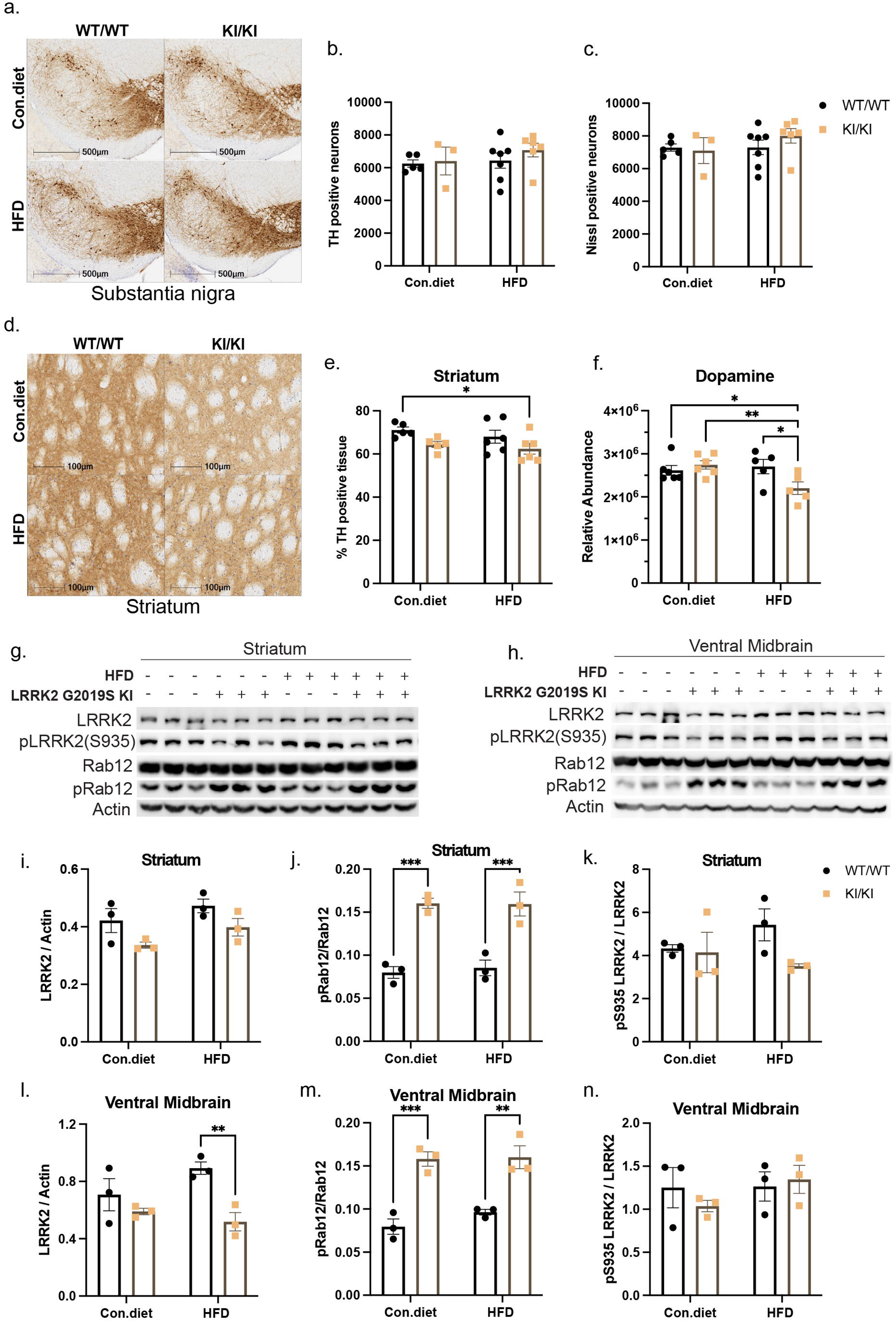
Pre-existing metabolic syndrome unmasks striatal dopamine loss without nigrostriatal dopaminergic neuronal loss in G2019S *LRRK2* knockin mice. a Representative image of TH immunostaining in the substantia nigra. Scale bar: 500 µm. Quantification of TH-positive neurons (**b**) and Nissl-positive neurons (**c**) in the substantia nigra (*n* = 3-7 per group). **d** Representative images of TH immunostaining in the striatum. Scale bar: 100 µm. **e** Quantification of TH-positive area in the striatum (*n* = 5-6 per group). **f** Relative striatal dopamine level measured by LC-MS (*n* = 5-6 per group). Western blot analysis (**g-h**) and quantification (**i-n**) of LRRK2 and Rab12 expression levels in the striatum (**g, i-k**) and ventral midbrain (**h, l-n**) of *G2019S LRRK2* knockin and wild-type mice under control (Con.) diet and HFD conditions (*n* = 3 per group). Data are presented as mean ± SEM. A two-way ANOVA was performed to evaluate the effects of high-fat diet and *G2019S LRRK2* mutant and their interaction followed by Fisher’s LSD comparisons. WT/WT, wild-type mice; KI/KI, homozygous *G2019S LRRK2* knockin mice. Data are presented as mean ± SEM. A two-way ANOVA was performed to evaluate the effects of high-fat diet and *G2019S LRRK2* mutant and their interaction followed by Fisher’s LSD comparisons. Significant main effects were found (\**P*<0.05, \*\**P*<0.01, \*\*\**P*<0.001).

LRRK2 expression varies across brain regions, with the highest levels observed in the striatum [15]. Key measures of LRRK2 kinase activity include its autophosphorylation at Serine 1292 (pS1292) and phosphorylation of its Rab substrates. To assess the effect of pre-existing MetS on LRRK2 kinase activity in mouse brain, we examined the levels of total and phosphorylated LRRK2 and Rab12 in striatal and ventral midbrain extracts (**Fig. 4 g-h**). We find a significant decrease of total LRRK2 levels in ventral midbrain of HFD-fed *G2019S LRRK2* knockin mice compared to WT mice, and a non-significant decrease in *G2019S LRRK2* knockin mice in general across brain regions relative to WT mice (**Fig. 4i, l**). Due to the suboptimal performance of the pS1292-LRRK2 antibody in brain tissue, we evaluated LRRK2 kinase activity solely by blotting for pRab12. As expected, we observe a significant ∼2-fold increase in pRab12 levels in *G2019S LRRK2* knockin mice compared to WT mice, but with no effect of HFD (**Fig. 4j, m**). Additionally, we do not observe any difference of pS935-LRRK2 levels between LRRK2 genotypes or diet, which serves as a useful marker for monitoring LRRK2 kinase inhibition *in vivo* (**Fig. 4o, r**). Our data demonstrate modestly reduced total LRRK2 levels and elevated LRRK2 substrate phosphorylation in the brains of *G2019S LRRK2* KI mice, but without a specific impact of diet.

### Altered pyrimidine metabolites in *G2019S LRRK2* knockin mice across multiple tissues

LRRK2 is a large multidomain protein with both GTPase and kinase activity, and is highly expressed in peripheral organs such as the lung and kidney [16]. It participates in a diverse set of cellular functions and signaling pathways including mitochondrial function, vesicle trafficking, endocytosis, retromer complex modulation and autophagy [16, 17]. However, the relationship between LRRK2 and metabolic pathways is poorly studied. To further elucidate the molecular mechanisms underlying *LRRK2*-linked PD and identify potential therapeutic targets, we assessed the effects of *G2019S LRRK2* on metabolic pathways in both neural and peripheral tissues, in the presence or absence of pre-existing MetS, using targeted metabolic profiling. As expected, HFD consumption induces metabolic alterations across multiple tissues (**Fig. S4**, **Table S1**), with the most pronounced changes observed in the striatum (**Fig. S4a, b**). Notably, we find that the pyrimidine nucleotides, thymidine and deoxyuridine, are significantly reduced in the striatum of *G2019S LRRK2* knockin mice compared to WT mice, under both control diet and HFD conditions (**Fig. 5a, b**), as well as in the plasma (**Fig 5c, d**) and lung (**Fig 5g, h**). In the liver, a significant reduction of thymidine and deoxyuridine is observed in *G2019S LRRK2* knockin mice under HFD condition (**Fig. 5f**) with a non-significant reduction under control diet (**Fig. 5e**). In the kidney, a significant reduction in thymidine levels and a non-significant decrease in deoxyuridine levels are observed in *G2019S LRRK2* knockin mice compared to WT mice, under both diets (**Fig. 5i, j**).

**Figure 5.**
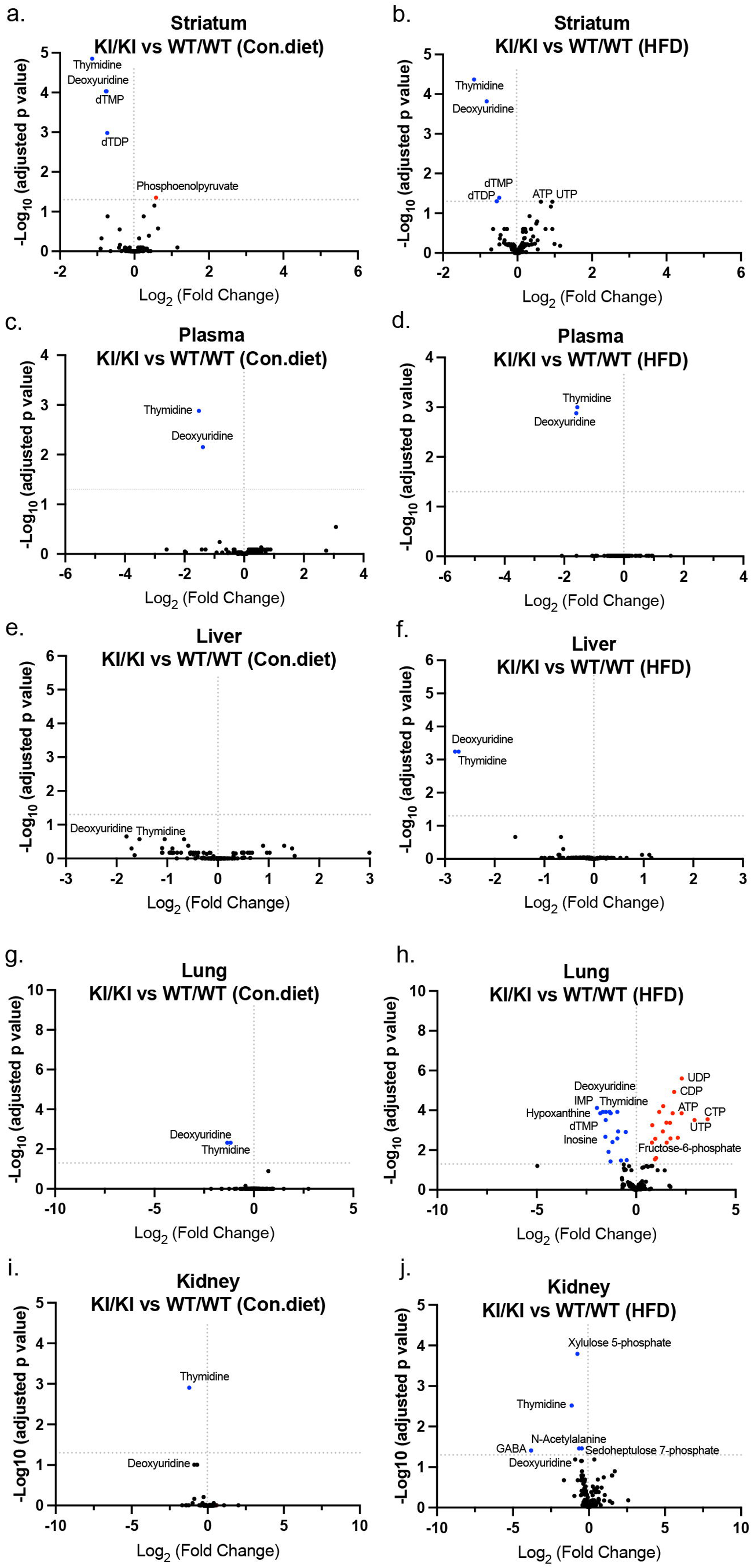
Non-lipid metabolite alterations across tissues in *G2019S LRRK2* knockin mice under Con. diet and HFD conditions. Volcano plot showing differences in the abundance of non-lipid metabolites between *G2019S LRRK2* knockin mice and wild-type controls under Con. diet and HFD in the striatum (**a, b**), plasma (**c-d**), liver (**e-f**), lung (**g-h)**, and kidney (**i-j**). WT/WT, wild-type mice; KI/KI, homozygous *G2019S LRRK2* knockin mice. Each point represents a non-lipid metabolite. X-axis shows log_2_ (fold change) in non-lipid metabolite abundance between *G2019S LRRK2* knockin mice and wild-type controls. Y-axis shows -log_10_ (adjusted *p*-value) from limma’s empirical Bayes moderated analysis (eBayes) combined with Benjamini-Hochberg (BH) for multiple testing correction (*n* = 6 per group). Non-lipid metabolites with adjusted *P*<0.05 are considered significantly altered. Significantly upregulated ones are highlighted in red and significantly downregulated ones are highlighted in blue.

To validate the observed alterations in pyrimidine metabolites in *G2019S LRRK2* knockin mice by targeted metabolic profiling, we performed absolute nucleoside quantitation in brain and plasma samples from 5-month-old *G2019S LRRK2* knockin and WT mice (**Fig. 6**). Consistent with our targeted metabolic profiling results, thymidine and deoxyuridine levels are markedly (>2-fold) and significantly reduced in *G2019S LRRK2* knockin mice compared to WT mice in the striatum (**Fig. 6a, b**), ventral midbrain (**Fig. 6f, g**), and plasma (**Fig. 6k, l**). Additionally, a modest increase in deoxycytidine levels is observed in the striatum of *G2019S LRRK2* knockin mice compared to WT mice, but not in ventral midbrain or plasma (**Fig. 6c, h, m**). The levels of deoxyadenosine and deoxyguanine are relatively low in brain yet undetectable in plasma, with no significant differences observed between genotypes (**Fig. 6d-e, i-j**).

**Figure 6.**
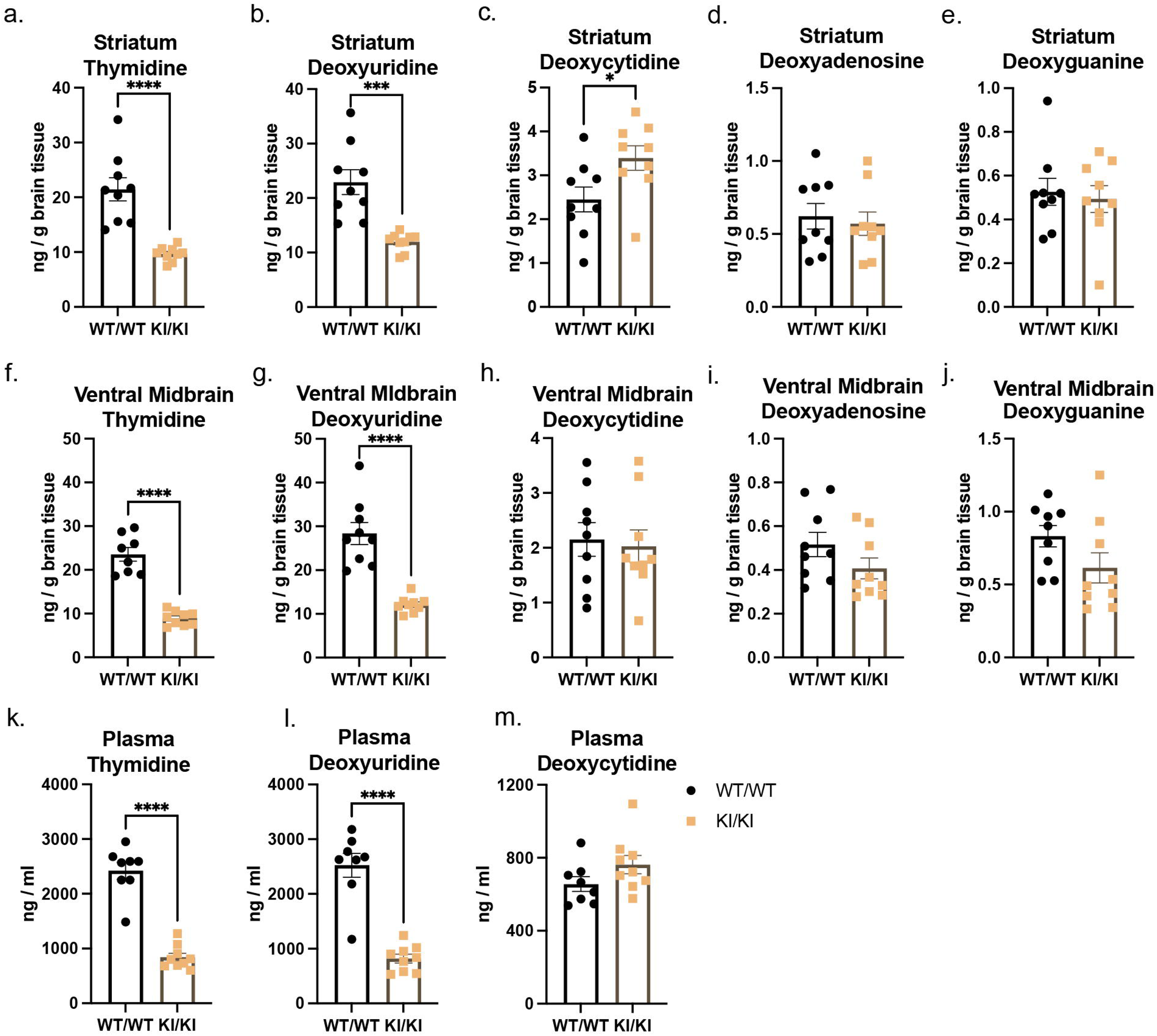
Absolute nucleoside quantitation in 5-month-old *G2019S LRRK2* knockin mice. Thymidine (**a**), deoxyuridine (**b**), deoxycytidine (**c**), deoxyadenosine (**d**), and deoxyguanine (**e**) concentrations in striatum (ng/g). Thymidine (**f**), deoxyuridine (**g**), deoxycytidine (**h**), deoxyadenosine (**i**), and deoxyguanine (**j**) concentrations in ventral midbrain (ng/g). Thymidine (**k**), deoxyuridine (**l**), and deoxycytidine (**m**) concentrations in plasma (ng/ml). Nucleoside quantity was measured by LC-MS (*n* = 8-9 per group). Data are presented as mean ± SEM. Unpaired two-tailed Student’s *t*-test was performed to evaluate the effects of *G2019S LRRK2* mutation (\**P*<0.05, \*\*\**P*<0.001, \*\*\*\**P*<0.0001).

### Pre-existing metabolic syndrome unmasks altered brain energy metabolism in *G2019S LRRK2* knockin mice

PD is increasingly recognized as a disorder involving disrupted brain energy homeostasis [18, 19]. As previously mentioned, we assessed the effects of *G2019S LRRK2* and pre-existing MetS on metabolic profiles in striatum, using targeted metabolic profiling (**Fig. 5a-b**, **S4a-b**). We find that the level of phosphoenolpyruvate and NADH is elevated in HFD-fed mice compared to control diet (**Fig. 7a-b**). Elevated levels of phosphoenolpyruvate (PEP) and NADH, alongside increased ATP (**Fig. 7c**) suggest that HFD consumption induces a glycolytic bottleneck and reduces ATP utilization in the brain. Interestingly, a synergistic effect of HFD and *G2019S LRRK2* is observed, resulting in the most pronounced ATP increase in HFD-fed *G2019S LRRK2* knockin mice (**Fig. 7c**), yet no difference in ATP levels between genotypes on control diet. Similar trends are observed in the levels of its related metabolites, ADP, UTP, and CTP (**Fig. 7d-f**). The increase of triphosphates and diphosphates (like ADP, UTP) indicates altered nucleotide turnover or salvage pathway upregulation. The metabolic alterations observed suggest a state of energy surplus in HFD-fed mice, characterized by increased energy production without efficient utilization, along with disrupted purine metabolism, particularly pronounced in *G2019S LRRK2* knockin mice.

**Figure 7.**
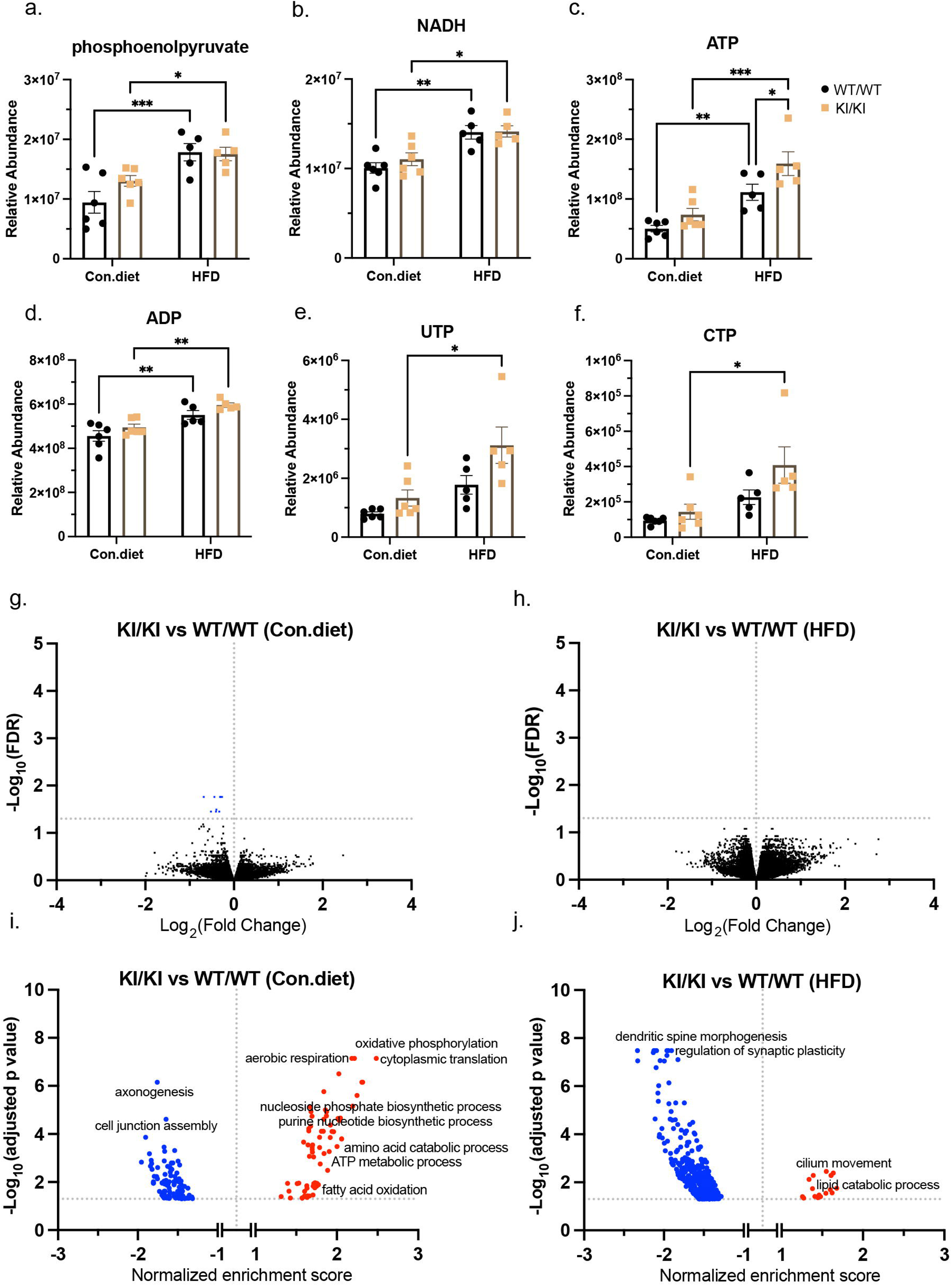
Pre-existing metabolic syndrome reveals altered brain energy metabolism in *G2019S LRRK2* knockin mice. **a-f** Bar plot showing comparation of selected metabolites (relative abundance) in the striatum across groups (*n* = 5-6 per group). These metabolites reflect alterations in energy metabolism (**a-c**) and nucleotide metabolism (**c-f**). Data are presented as mean ± SEM. A two-way ANOVA was performed to evaluate the effects of high-fat diet and *G2019S LRRK2* mutant and their interaction, Tukey’s post-hoc test was used for multiple comparisons. Significant main and interaction effects were found (\**P*<0.05, \*\**P*<0.01, \*\*\**P*<0.001). **g-h** Volcano plot showing differential gene expression in the striatum between *G2019S LRRK2* knockin and wild-type mice under Con.diet and HFD conditions. Each point represents one gene. X-axis shows log_2_ (fold change) in gene expression, Y-axis shows -log_10_ (adjusted *p*-value), significance was defined as a Benjamini–Hochberg adjusted *P*<0.05 (*n* = 4 per group). Gene expression with adjusted *P*<0.05 are considered significantly altered. Significantly downregulated genes are highlighted in blue. **i-j** Volcano plot illustrates enriched biological processes between *G2019S LRRK2* knockin and wild-type mice under Con.diet and HFD conditions as identified by gene set enrichment analysis (GSEA). Each point represents one process, X-axis shows normalized enrichment score, Y-axis shows -log_10_ (adjusted *p*-value), significance was defined as a Benjamini–Hochberg adjusted *P*<0.05 (*n* = 4 per group). Significantly upregulated genes are highlighted in red and significantly downregulated genes are highlighted in blue. WT/WT, wild-type mice; KI/KI, homozygous *G2019S LRRK2* knockin mice.

To gain further insight into the metabolic alterations observed in the striatum of *G2019S LRRK2* knockin mice, we performed bulk RNA sequencing of striatal tissue. We find that HFD consumption markedly altered the gene expression profile in the striatum of both WT and *G2019S LRRK2* knockin mice, with many genes showing differential expression (**Fig. S5a, b**, **Table S2**). However, only a limited number of genes are differentially expressed between *G2019S LRRK2* knockin and WT mice under either dietary condition (**Fig. 7g-h**). Gene set enrichment analysis reveals that, compared to control diet-fed mice, HFD exposure markedly alters the biological processes in the striatum of both WT and *G2019S LRRK2* knockin mice (**Fig. S5c-d**). Interestingly, compared to control diet-fed WT mice, *G2019S LRRK2* knockin mice exhibit the upregulation of similar striatal biological processes to those induced by HFD (**Fig. 7i**, **S5c**, **Table S3**). Notably, oxidative phosphorylation and purine nucleoside biosynthetic processes are elevated in *G2019S LRRK2* knockin mice compared to WT mice under control diet conditions (**Fig. 7i**). Moreover, under HFD conditions, most of these enriched processes are oppositely downregulated in *G2019S LRRK2* knockin mice compared to WT controls (**Fig. 7j**), suggesting a potential suppression of cellular activity in *G2019S LRRK2* knockin mice along with a reduction in cellular energy demand or utilization. Together, the transcriptomic alterations observed by RNA-seq analysis align with and help explain the metabolite changes identified in the striatum, supporting a coordinated disruption of energy and nucleotide metabolism in *G2019S LRRK2* knockin mice, particularly under HFD conditions.

We next evaluated whether similar or complementary transcriptional changes occurred in the ventral midbrain. We performed single-nucleus RNA sequencing (snRNA-seq) on ventral midbrain tissues from *G2019S LRRK2* knockin and WT mice under control diet and HFD to uncover cell type-specific transcriptional responses to *G2019S LRRK2* and metabolic stress. A total of 34,969 nuclei were recovered and grouped into 27 transcriptionally distinct cell clusters (**Fig. 8a**). Using established cell-type markers (**Fig. S6a**), each cluster was annotated, revealing major neural and glial populations, along with a small subset of TH-positive dopaminergic neurons grouped in cluster 27 (**Fig. 8a**, **S8b**). Oligodendrocytes comprise the largest cell population in the ventral midbrain, followed by neurons and astrocytes (**Fig. S6b**). This unique dataset enables the dissection of cell type-specific transcriptional alterations driven by genetic and metabolic factors. Visualization of LRRK2 expression across cell clusters reveals highest expression in astrocytes (cluster 1), oligodendrocytes (cluster 2 and 22), and fibroblasts (cluster 3, 7 and 22) (**Fig. 8b**). Surprisingly, we find that only a small number of genes are differentially expressed between groups and across cell types (**Table S4**). Consistent with our bulk RNA-seq data, gene set enrichment analysis reveals that most biological processes that are enriched in *G2019S LRRK2* KI mice relative to WT mice on a control diet, are now oppositely downregulated by exposure to a HFD diet across multiple cell types, suggesting a potential suppression of cellular activity (**Table S5**). Notably, oxidative phosphorylation is significantly downregulated in astrocytes and oligodendrocytes in HFD-fed *G2019S LRRK2* knockin mice (**Fig. 8c**, **Table S5**) yet upregulated in these mice on a control diet. Moreover, consistent with bulk RNA-seq data, *G2019S LRRK2* knockin mice on a control diet exhibit a similar non-significant upregulation of oxidative phosphorylation across cell types to those induced by HFD in WT mice (**Fig. 8c**). These findings suggest that metabolic stress unmasks a cell type-specific (i.e. astrocytes and oligodendrocytes) vulnerability in brain energy metabolism associated with G2019S LRRK2. Additionally, gene set enrichment analysis reveals that, under HFD conditions, the response to insulin and insulin receptor signaling pathway are downregulated in *G2019S LRRK2* knockin mice compared to WT mice in neurons, which could potentially lead to impaired neuronal function, altered synaptic transmission, and neurodegenerative effects (**Table S5**). Overall, these data suggest that G2019S LRRK2 and HFD exposure induce similar transcriptional responses in the mouse ventral midbrain, particularly oxidative phosphorylation genes across cell types. Furthermore, the normal transcriptional response of WT mice to HFD exposure is impaired in *G2019S LRRK2* mice.

**Figure 8.**
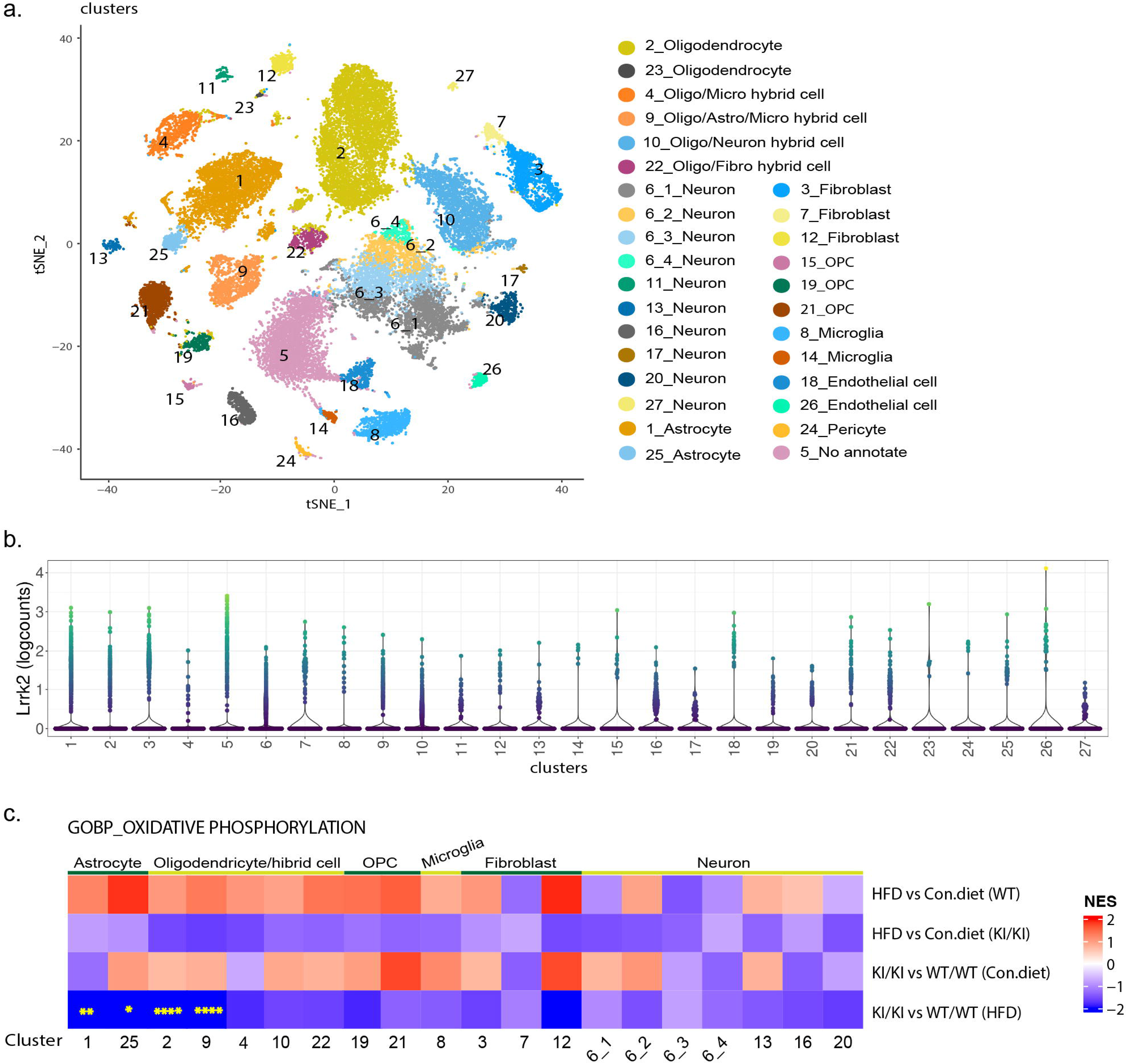
Cell type-specific transcriptional responses to *G2019S LRRK2* and metabolic stress. Nuclei were isolated from ventral midbrain of *G2019S LRRK2* knockin mice and their littermates (*n* = 4 per group), 34,969 nuclei were recovered in total. **a** *t*-SNE map shows 27 classified cell clusters with annotations. **b** Violin plot shows Lrrk2 expression across clusters. **c** Heatmap illustrates oxidative phosphorylation pathway enrichment across multiple cell types in various group comparisons. Each square represents a specific comparison (labeled on the right) within a given cell cluster (labeled at the bottom). Red squares indicate positive enrichment scores; blue squares indicate negative enrichment scores. Comparisons with statistically significant (Benjamini–Hochberg adjusted *p*) enrichment are marked with stars (\**P*<0.05, \*\**P*<0.01, \*\*\*\**P*<0.0001). WT/WT, wild-type mice; KI/KI, homozygous *G2019S LRRK2* knockin mice.

### Thymidine phosphorylase is upregulated in the liver of *G2019S LRRK2* knockin mice

The observed alterations in pyrimidine metabolites in *G2019S LRRK2* knockin mice prompted us to examine enzymes involved in its metabolism. Thymidine phosphorylase (TP), encoded by the *TYMP* gene, catalyzes the reversible phosphorolysis of thymidine and deoxyuridine into their respective bases (thymine and uracil) and 2-deoxyribose 1-phosphate, primarily functioning in a catabolic direction (**Fig. 9a**) [20–22]. Given its central role in pyrimidine metabolism and its predominant expression in the liver in mice [23], we first examined thymidine phosphorylase levels in liver and brain. We find that thymidine phosphorylase protein levels are significantly elevated in the liver, but not obviously in the brain, of *G2019S LRRK2* knockin compared to WT mice (**Fig. 9b**). Notably, TP protein is highly abundant in the liver relative to brain. To further investigate the molecular basis of the altered pyrimidine metabolites in these mice, we performed qRT-PCR on brain tissue and bulk RNA sequencing on liver tissue from *G2019S LRRK2* knockin and WT mice. Consistent with increased TP protein, *TYMP* mRNA is significantly upregulated in the liver (**Fig. 9d**), and is also non-significantly increased by ∼2-fold in the brain (**Fig. 9c**), of *G2019S LRRK2* knockin mice compared to WT mice. In the liver, *TYMP* represents the largest differentially expressed upregulated gene in *G2019S LRRK2* KI mice (**Fig. 9d**). Gene set enrichment analysis reveals that processes related to fatty acid and lipid metabolism are upregulated in the liver of *G2019S LRRK2* knockin mice compared to WT mice (**Fig. 9e**). To ensure the reproducibility of TP protein changes and to evaluate the combined effects of HFD and G2019S LRRK2 on its activity in the liver, we performed Western blot analysis in the HFD-fed mouse cohort (**Fig. 9f**). We find a similar significant increase of thymidine phosphorylase protein in *G2019S LRRK2* knockin mice irrespective of diet (**Fig. 9g**), suggesting that TP upregulation in the liver is primarily driven by G2019S LRRK2. Total LRRK2 protein levels exhibit inter-individual variability among mice (**Fig. 9h**), without being significantly different between groups, which may be attributable to the relatively low levels of LRRK2 protein detected in liver compared with brain and lung (**Fig. S7**). Under control diet conditions, male mice appear to exhibit higher LRRK2 protein levels than female mice, however, the small and unbalanced sample size (2 males vs. 4 females) limits interpretation of potential sex-related differences. We do not observe differences in the levels of pSer935-LRRK2 (**Fig. 9j**). The level of pSer1292-LRRK2 is increased in *G2019S LRRK2* knockin mice compared to WT mice with no consistent effect of control diet or HFD (**Fig. 9i**). We further assessed LRRK2 kinase activity by blotting for its substrates pRab10 and pRab12 (**Fig. 9f**). The level of pRab10 and pRab12 are elevated in *G2019S LRRK2* knockin mice compared with WT mice (**Fig. 9l-m**), with no significant differences observed between HFD and control diet groups. These data demonstrate that *TYMP* mRNA and TP protein are markedly increased in the liver (and brain) of *G2019S LRRK2* KI mice, most likely accounting for the systemic decrease in thymidine and deoxyuridine levels. However, HFD exposure does not consistently alter LRRK2 kinase activity in the liver.

**Figure 9.**
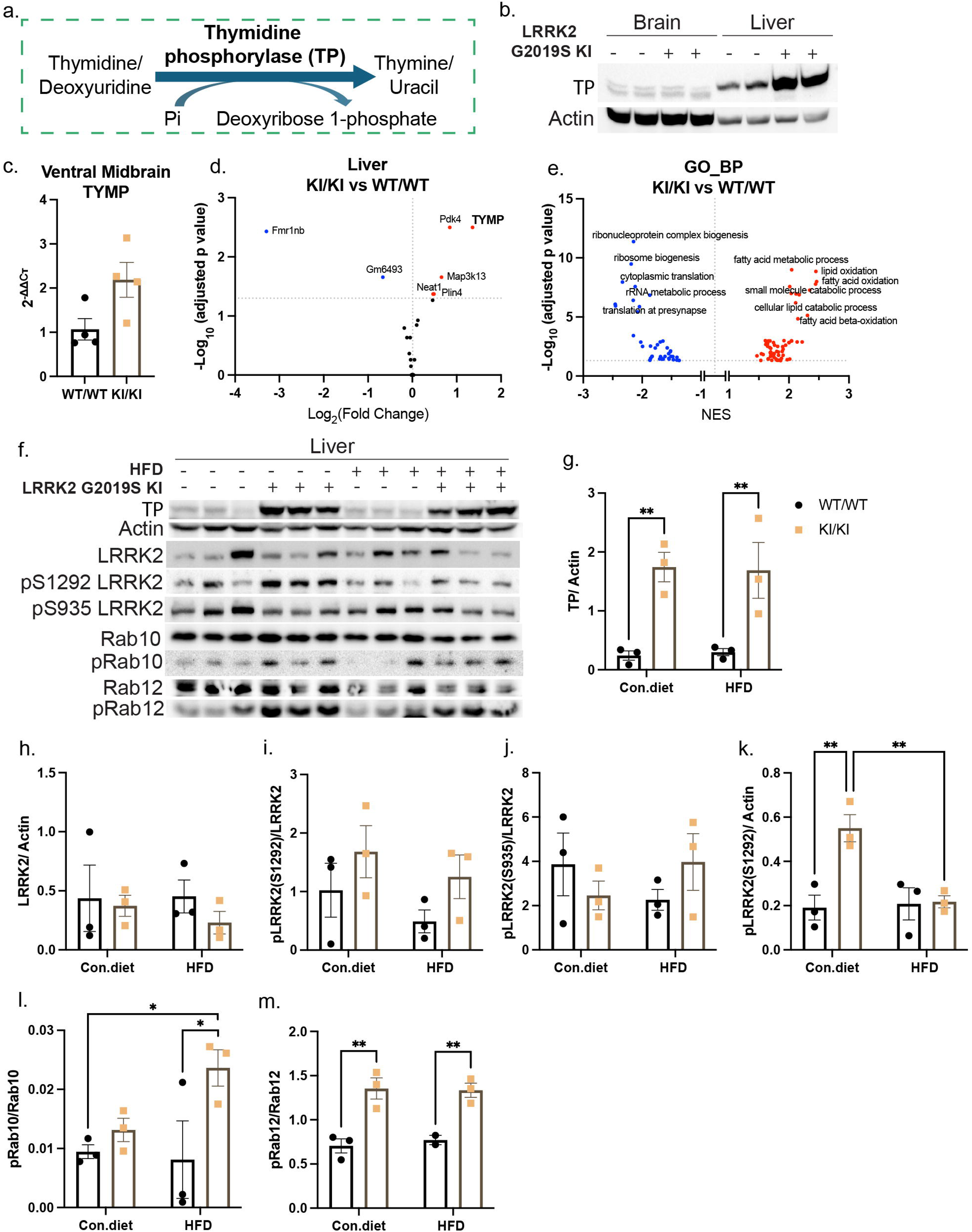
Thymidine phosphorylase is upregulated in the liver of *G2019S LRRK2* knockin mice. **a** Diagram showing that thymidine phosphorylase catalyzes the conversion of thymidine and deoxyuridine into their respective bases (thymine and uracil) and 2-deoxyribose 1-phosphate. **b** Western blot analysis of thymidine phosphorylase (TP) expression levels in brain and liver in 5-month-old *G2019S LRRK2* knockin mice and controls (*n* = 2 per group). **c** qRT-PCR analysis of *TYMP* levels in ventral midbrain of 5-month-old *G2019S LRRK2* knockin mice and controls (*n* = 4 per group). Data are presented as mean ± SEM. Unpaired two-tailed Student’s *t*-test was performed to evaluate the effects of *G2019S LRRK2* mutation, no significant effect was found. **d** Volcano plot showing differential gene expression in the liver of 5-month-old G2019S LRRK2 knockin mice compared to wild-type controls. Each point represents one gene. X-axis shows log_2_ (fold change) in gene expression in *G2019S LRRK2* knockin mice compared to wild-type controls. Y-axis shows -log_10_ (adjusted *p*-value) from limma’s empirical Bayes moderated analysis (eBayes) combined with Benjamini-Hochberg (BH) for multiple testing correction (*n* = 4 per group). Gene expression with adjusted *P*<0.05 are considered significantly altered. Significantly upregulated genes are highlighted in red and significantly downregulated genes are highlighted in blue. **e** Volcano plot illustrating enriched biological processes in *G2019S LRRK2* knockin mice compared to wild-type controls as identified by gene set enrichment analysis (GSEA). Each point represents one process. X-axis shows normalized enrichment score, Y-axis shows -log_10_ (adjusted *p*-value), significance was defined as a Benjamini–Hochberg adjusted *P*<0.05 (*n* = 4 per group). Significantly upregulated pathways are highlighted in red and significantly downregulated pathways are highlighted in blue. **f** Western blot analysis and quantification (**g-m**) of TP, total and phosphorylated LRRK2, Rab10, and Rab12 expression levels in the liver of *G2019S LRRK2* knockin and wild-type mice under Con.diet and HFD conditions (*n* = 3 per group). WT/WT, wild-type mice; KI/KI, homozygous *G2019S LRRK2* knockin mice. Data are presented as mean ± SEM. A two-way ANOVA was performed to evaluate the effects of high-fat diet and *G2019S LRRK2* mutant and their interaction followed by Fisher’s LSD comparisons. Significant main effects were found (\**P*<0.05, \*\**P*<0.01).

### Pre-existing metabolic syndrome and G2019S LRRK2 synergistically disrupt central carbon and nucleotide metabolism in the lung

LRRK2 is highly expressed in lung, and its metabolic state may reveal peripheral consequences of LRRK2 dysregulation, offering insights into potential pulmonary effects of LRRK2 inhibitor therapies. As previously mentioned, we assessed the effects of *G2019S LRRK2* and pre-existing MetS on the metabolic profile of lung tissue using targeted metabolic profiling (**Fig. 5g-h**, **S4g-h**). We find that glucose-6-phosphate (G-6-P) and fructose-6-phosphate (F-6-P) are significantly increased in *G2019S LRRK2* knockin mice under HFD conditions compared to controls (**Fig. 10a-c**), suggesting that a bottleneck exists in glycolysis in the lung. Glucose is being phosphorylated but not efficiently processed downstream, leading to metabolite buildup (G-6-P, F-6-P) and most likely depletion downstream. The levels of NADH and ATP are significantly increased in HFD-fed *G2019S LRRK2* knockin mice compared to HFD-fed WT mice or control diet-fed *G2019S LRRK2* knockin mice (**Fig. 10d, n**), reflecting enhanced energy production or reduced consumption in the lung. Increased 2-ketoglutarate (2-KG) levels in HFD-fed *G2019S LRRK2* knockin mice suggests citric acid (TCA) cycle flux may be altered, possibly stalled after 2-KG, contributing to metabolite accumulation (**Fig. 10e**). Direct metabolic flux analysis would be required to further conclusively demonstrate altered TCA cycle flux. Additionally, we find increased NADPH levels and decreased xylulose-5-phosphate levels in HFD-fed *G2019S LRRK2* knockin mice compared to HFD-fed WT mice or control diet-fed *G2019S LRRK2* knockin mice (**Fig. 10f-g**). Increased NADPH suggests activation of the oxidative Pentose Phosphate Pathway (PPP), but the decrease in xylulose-5-phosphate (a non-oxidative PPP intermediate) indicates suppression of the non-oxidative branch, potentially impairing nucleotide biosynthesis. In addition to the interaction between the *G2019S LRRK2* and HFD affecting metabolite levels, we also observe that either *G2019S LRRK2* or HFD alone is sufficient to reduce xylulose-5-phosphate levels (**Fig. 10g**), indicating *G2019S LRRK2* exerts a similar suppressive effect on the non-oxidative branch of the PPP as HFD. Similarly, both *G2019S LRRK2* and HFD independently reduce IMP levels, with a robust additive effect observed in HFD-fed *G2019S LRRK2* knockin mice (**Fig. 10h**). This additive effect might suggest that G2019S LRRK2 and HFD do not lower IMP levels through the same pathway or mechanism. Consistent with the reduction in IMP, similar trends are observed for the levels of its related metabolites, inosine and hypoxanthine, culminating in a robust reduction in HFD-fed *G2019S LRRK2* knockin mice (**Fig. 10i-j**). Similar to increased ATP levels, we also observe a marked increase of GDP, GTP, ADP, UTP, and CTP levels in HFD-fed *G2019S LRRK2* knockin mice compared to HFD-fed WT mice or control diet-fed *G2019S LRRK2* knockin mice (**Fig. 10k-p**). Notably, these nucleotides are not different between *G2019S LRRK2* knockin and WT mice at baseline on a control diet, or between WT mice on a control diet or HFD (**Fig. 10k-p**), implying that the combination of G2019S LRRK2 and HFD drives these increases. The reduction in IMP suggests impaired *de novo* purine synthesis, possibly due to a lack of ribose precursors owing to suppressed non-oxidative PPP. The increase in downstream triphosphates and diphosphates (like GTP, ADP) indicates altered nucleotide turnover or salvage pathway upregulation. The metabolic alterations observed here reveal that *G2019S LRRK2*, especially under HFD exposure, leads to a coordinated disruption of glycolysis, the pentose phosphate pathway, and nucleotide metabolism in the lung.

**Figure 10.**
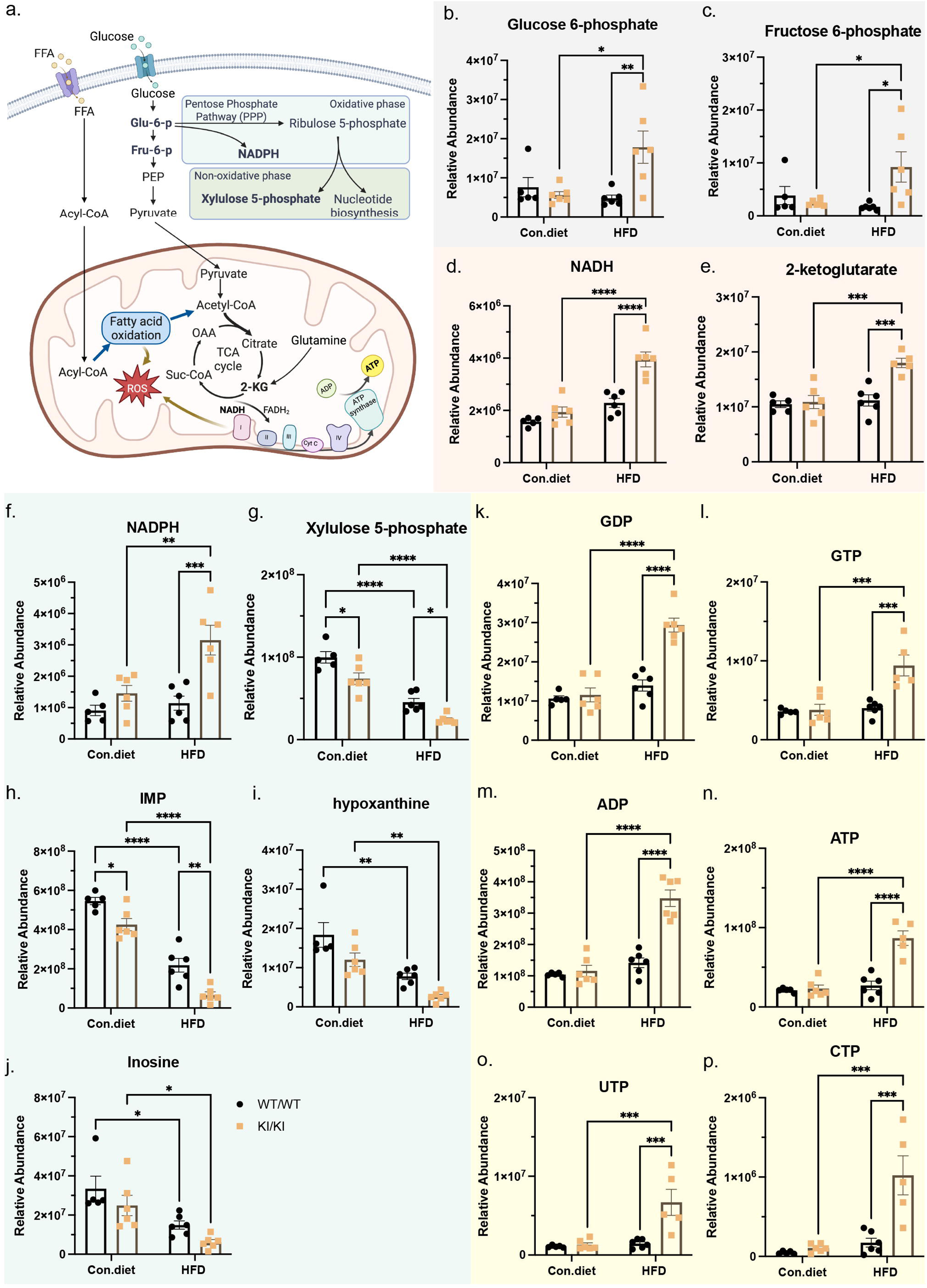
Pre-existing metabolic syndrome and *G2019S LRRK2* synergistically disrupt lung metabolism. **a** Schematic diagram showing central energy and biosynthetic pathways involving glucose and fatty acids. **b-p** Bar plot showing comparison of selected metabolites (relative abundance) in lung tissue from *G2019S LRRK2* knockin and control mice exposed to either HFD or control diet (*n* = 6 mice/group). These metabolites reflect alterations in glycolysis (**b-c**), TCA cycle (**d-e**), pentose phosphate pathway (**f-j**), and nucleotide metabolism (**k-p**). Data are presented as mean ± SEM in bar plot. A two-way ANOVA was performed to evaluate the effects of high-fat diet and *G2019S LRRK2* and their interaction, Tukey’s post hoc test was used for multiple comparisons. Significant main and interaction effects were found (\**P*<0.05, \*\**P*<0.01, \*\*\**P*<0.001, \*\*\*\**P*<0.0001).

HFD consumption has been shown to induce inflammation and increase the risk of lung fibrosis in mice [24, 25]. To assess the impact of pre-existing metabolic syndrome on lung pathology in *G2019S LRRK2* knockin mice, we performed histological analysis of lung tissue using H&E staining (**Fig. 11a**). Increased inflammatory cell infiltration around the airways and blood vessels is consistently observed in *G2019S LRRK2* knockin and WT mice under HFD conditions (**Fig. 11a**). To validate the observed immune cell infiltration, we performed immunostaining of lung using the macrophage marker CD68 (**Fig. 11b**) and T cell marker CD3 (**Fig. 11c**). Consistent with H&E staining, we observe an increased number of CD68-positive and CD3-positive cells surrounding the airways and blood vessels in HFD-fed mice (**Fig. 11b-c**), indicating the presence of an inflammatory environment in the lung. This HFD-driven inflammation may contribute to the metabolic alterations observed in the lungs of *G2019S LRRK2* knockin mice. To further explore these metabolic alterations, we conducted proteomic analysis of lung tissues. Comparison of total proteome profiles reveals marked alterations in HFD-fed mice relative to control diet-fed mice, indicating that HFD consumption significantly reshapes the lung proteome (**Fig. 11d-e**, **Table S6**). However, no significant differences in the total proteome are detected between *G2019S LRRK2* knockin and WT mice under either dietary condition (**Fig. 11f-g**), suggesting that protein post-translational modifications may instead contribute to the metabolic alterations observed in the lungs of *G2019S LRRK2* knockin mice. To assess the effects of HFD and the G2019S mutation on LRRK2 kinase activity in the lung, we performed Western blot analysis of total and phosphorylated LRRK2 (**Fig. 11h**). We find that pSer1292-LRRK2 levels are increased in *G2019S LRRK2* knockin mice compared to WT mice (**Fig. 11j, l**), whereas pSer935-LRRK2 levels remain unchanged among groups (**Fig. 11k**). Interestingly, total LRRK2 levels are decreased in *G2019S LRRK2* knockin mice compared to WT mice irrespective of diet (**Fig. 11i**), similar to brain LRRK2 (**Fig. 4**), suggesting an unappreciated effect of the G2019S mutation on the protein stability of endogenous LRRK2. We next compared the levels of phosphorylated Rab8a, Rab10, and Rab12 among groups (**Fig. 11h**). The levels of pRab8a, pRab10, and pRab12 are all increased in *G2019S LRRK2* knockin mice compared to WT mice under HFD conditions, and either show a significant increase (pRab8a) or a non-significant increase (pRab10, pRab12) under control diet conditions (**Fig. 11m-o**). These data suggest that LRRK2-mediated Rab phosphorylation (and pSer1292-LRRK2) is enhanced by the G2019S mutation, as expected, yet with no obvious effect of HFD alone or when combined. Overall, our data suggest that post-translational regulation of LRRK2, or downstream pathways, may contribute to the metabolic alterations observed in the lungs of *G2019S LRRK2* knockin mice.

**Figure 11.**
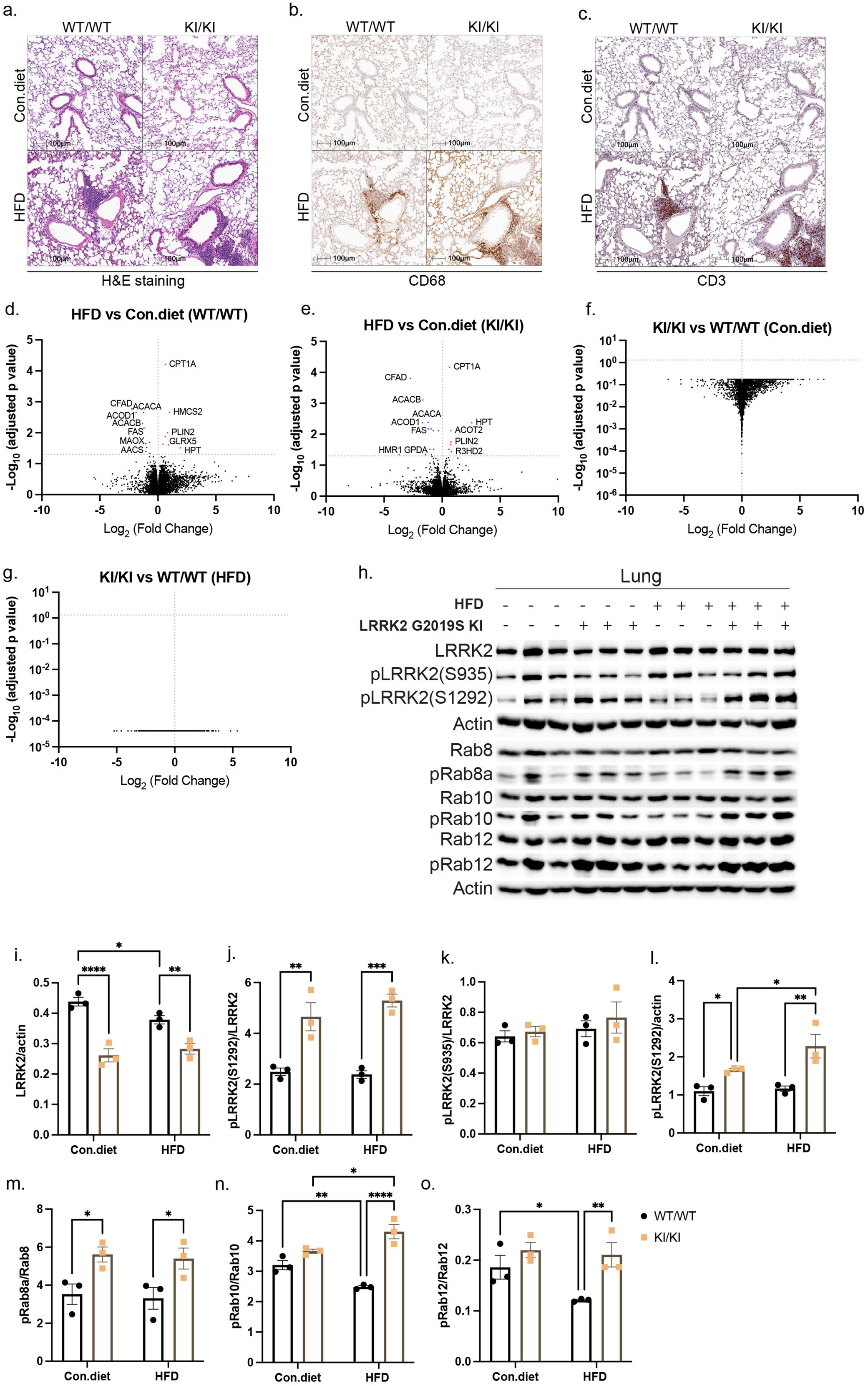
Pre-existing metabolic syndrome increases inflammation and reshapes the lung proteome in *G2019S LRRK2* knockin mice. **a-c** Representative images of H&E staining (**a**), CD68 (**b**) and CD3 (**c**) immunostaining of the lungs. Scale bar: 100 µm. **d-g** Volcano plot showing differences in the abundance of proteins between HFD and Con. diet groups in wild-type mice (**d**) and *G2019S LRRK2* knockin mice (**e**); between *G2019S LRRK2* knockin and wild-type mice under Con. diet (**f**) and HFD (**g**) conditions (*n* = 4 mice/group). Each point represents a protein. X-axis shows log_2_ fold change in protein abundance between each comparison. Y-axis shows -log_10_ adjusted *p*-value from limma’s empirical Bayes moderated analysis (eBayes) combined with Benjamini-Hochberg (BH) for multiple testing correction (*n* = 4 mice/group). Proteins with adjusted *P*<0.05 are considered significantly altered. Significantly upregulated proteins are highlighted in red and significantly downregulated proteins are highlighted in blue. **h** Western blot analysis and quantification (**i-o**) of total and phosphorated LRRK2, Rab8a, Rab10 and Rab12 expression levels in the lung of *G2019S LRRK2* knockin and wild-type mice under Con.diet and HFD conditions (*n* = 3 mice/group). WT/WT, wild-type mice; KI/KI, homozygous *G2019S LRRK2* knockin mice. Data are presented as mean ± SEM. A two-way ANOVA was performed to evaluate the effects of high-fat diet and *G2019S LRRK2* and their interaction by Fisher’s LSD comparisons. Significant main and interaction effects were found (\**P*<0.05, \*\**P*<0.01, \*\*\**P*<0.001, \*\*\*\**P*<0.0001).

## Discussion

PD is recognized as a multifactorial disorder resulting from complex interactions among genetic risk, environmental exposure, aging and other factors [26]. The *G2019S LRRK2* mutation is the most common cause of monogenic PD and is also associated with an increased risk of sporadic PD [27]. However, penetrance of the *G2019S* mutation is incomplete, indicating that additional factors contribute to disease onset and progression [3, 27]. HFD is a well-known environmental/metabolic stressor linked to insulin resistance, mitochondrial dysfunction, and neuroinflammation, which is also implicated in PD [28, 29]. Combining HFD with LRRK2 PD models helps reveal whether metabolic stress unmasks or accelerates PD phenotypes in a susceptible genetic background. In this study, we investigate the interaction between the *G2019S LRRK2* mutation and chronic metabolic stress induced by prolonged HFD feeding in aged mice, using a multi-organ, multi-omics approach, including metabolomics, proteomics, bulk RNA sequencing, and single-nucleus RNA sequencing. We identify coordinated metabolic disturbances in the brain, liver, lung and kidney in *G2019S LRRK2* knockin mice. Our findings demonstrate that pre-existing metabolic syndrome unmasks widespread disruptions in systemic nucleotide and energy metabolism and exacerbates mitochondrial dysfunction in *G2019S LRRK2* knockin mice.

Metabolomics profiling revealed that thymidine and deoxyuridine levels are consistently and robustly reduced across distinct tissues in *G2019S LRRK2* knockin mice, most likely due to increased hepatic expression of thymidine phosphorylase (TP). TP catalyzes the catabolism of pyrimidine nucleosides, and its upregulation could therefore explain their systemic depletion [20]. Since these two nucleosides are essential for mitochondrial DNA replication and repair, their depletion may compromise mitochondrial integrity, a process highly relevant to PD pathogenesis [30, 31]. Notably, differences exist between human and mouse TP biology [23]. In humans, TP is broadly expressed in multiple tissues, including platelets, endothelial cells, and liver, and functions as an angiogenic factor (PD-ECGF) [21]. By contrast, in mice, TP expression is more restricted, with the liver serving as a primary site of activity [23]. Therefore, the observed hepatic TP upregulation in *G2019S LRRK2* knockin mice may have disproportionately strong effects on systemic nucleoside metabolism compared with humans. These alterations are likely to reflect secondary or compensatory adaptations in pyrimidine metabolism, potentially triggered by systemic stress responses or liver-specific metabolic vulnerability. Given the liver-enriched expression of TP in mice, it is possible that *LRRK2* mutations perturb hepatic homeostasis in a manner that converges on nucleoside metabolism, even though the mechanism may not be directly tied to canonical LRRK2 activity. These findings emphasize the need to consider systemic and tissue-specific effects of LRRK2 perturbations when interpreting metabolic phenotypes.

Alterations in purine and energy metabolism were not apparent in *G2019S LRRK2* knockin mice under control diet conditions but became evident upon exposure to a high-fat diet in lung and brain. Specifically, ATP levels were elevated significantly in *G2019S LRRK2* mice in the context of HFD, suggesting that the G2019S LRRK2 mutation creates a latent susceptibility in nucleotide and energy homeostasis that requires metabolic stress to manifest. This conditional “two-hit” phenotype highlights the importance of environmental factors, such as diet, in unmasking metabolic vulnerabilities in PD-linked genetic backgrounds.

Single-nucleus RNA sequencing analysis reveals that astrocytes and oligodendrocytes in the ventral midbrain are the most disrupted in oxidative phosphorylation transcriptional pathways in *G2019S LRRK2* knockin mice under HFD conditions. As key providers of metabolic and trophic support, glial dysfunction is likely to have downstream consequences for dopaminergic neurons [32–35]. Astrocytes regulate extracellular nucleotide turnover and shuttle energy substrates such as lactate to neurons, while oligodendrocytes sustain axonal energy needs through mitochondrial and glycolytic support [33, 35]. Impairment of these glial populations could compromise the bioenergetic environment necessary for dopaminergic neuron survival and synaptic function [33, 35]. In PD, dopaminergic neurons are highly energy-demanding and particularly sensitive to mitochondrial dysfunction [36]. The observation that glial oxidative phosphorylation is preferentially disrupted in *G2019S LRRK2* mice under HFD suggests a non-cell-autonomous mechanism whereby systemic purine imbalance and glial energy failure could eventually exacerbate dopaminergic neuronal vulnerability. This is consistent with human metabolomic studies reporting purine alterations (hypoxanthine, inosine) and evidence of impaired mitochondrial metabolism in serum or CSF of PD subjects [37–42]. The convergence of our findings in *G2019S LRRK2* mice with human biospecimen data supports the idea that purine metabolites in serum/CSF, together with markers of glial mitochondrial dysfunction, may serve as candidate biomarkers of *LRRK2*-PD progression. Moreover, these disrupted metabolites and pathways may provide therapeutic entry points to stabilize metabolic homeostasis and potentially protect neuronal function in individuals genetically at-risk of developing PD.

In this study, the differing magnitude of phenotypes observed between peripheral tissues and the brain may be influenced by tissue-specific LRRK2 expression levels (**Fig. S7**) as well as differences in the response of distinct tissues to secondary stressors, such as HFD-induced metabolic or inflammatory burden (**Fig. 12**). To further strengthen the observed metabolic changes in this “two-hit” mouse model, additional functional studies are now warranted and could prove informative. For example, i) metabolic cage analysis could assess whole-body energy expenditure and substrate utilization, ii) Seahorse analysis could evaluate mitochondrial respiration and bioenergetic capacity in different tissues, and iii) stable isotope-labeled substrate tracing could directly quantify metabolic flux through key energy pathways in different tissues. Such functional metabolic assays will form the basis of future studies.

**Figure 12.**
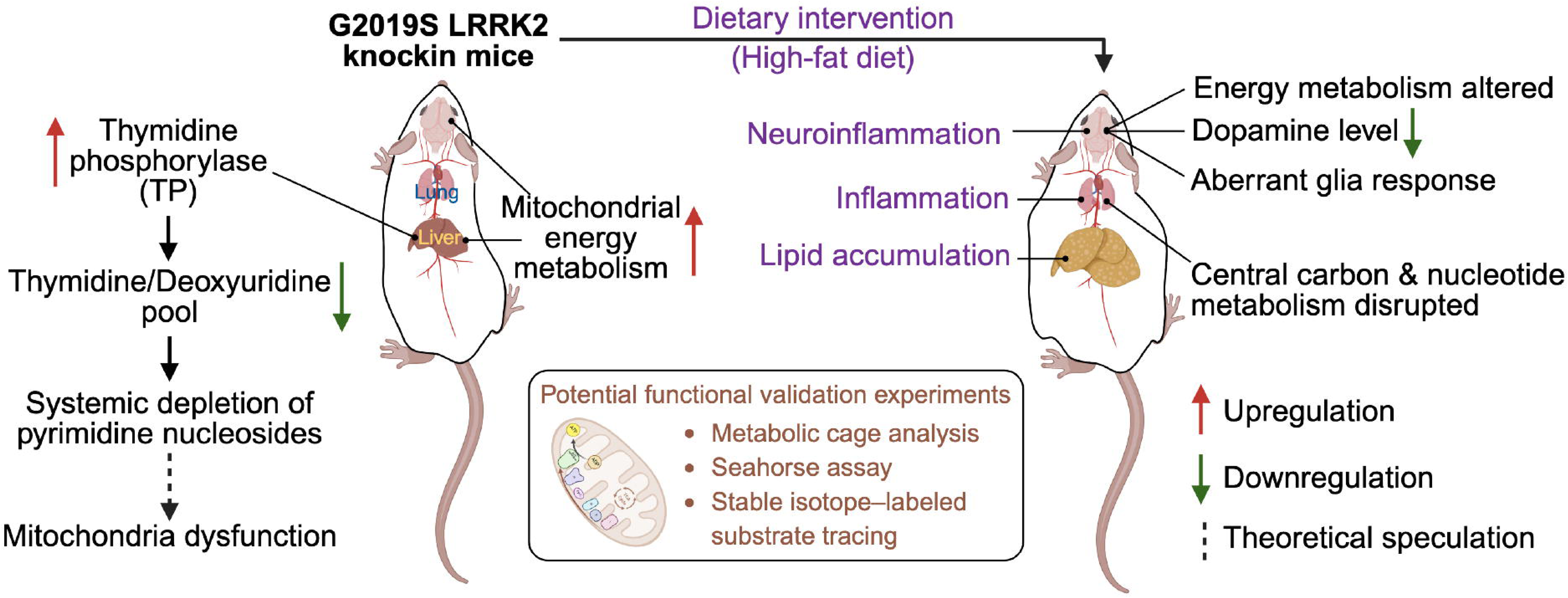
Graphical summary. Thymidine phosphorylase levels were significantly increased in the liver of *G2019S LRRK2* knockin mice, leading to systemic depletion of pyrimidine nucleosides, which may contribute to mitochondrial dysfunction. Under control diet conditions, increased mitochondrial energy metabolism was observed in both brain and peripheral tissues of *G2019S LRRK2* knockin mice, potentially reflecting a compensatory response to cellular stress or damage. High-fat diet exposure induced systemic metabolic dysregulation in *G2019S LRRK2* knockin mice, accompanied by hepatic lipid accumulation, pulmonary inflammation, and neuroinflammatory responses in the brain. The combined effects of inflammation and the *G2019S* mutation revealed disruption of central carbon and nucleotide metabolism in the lung, as well as alterations in energy metabolism, decreased dopamine levels, and aberrant glial responses in the brain. Potential functional validation experiments, including metabolic cage analysis, Seahorse mitochondrial respiration assays, and stable isotope-labeled substrate tracing, would be informative.

In conclusion, our findings demonstrate that pre-existing metabolic syndrome, induced by chronic high-fat diet feeding, unmasks widespread disruptions in systemic nucleotide and energy metabolism and exacerbates mitochondrial dysfunction in *G2019S LRRK2* knockin mice. Importantly, many of these effects are not evident under basal conditions, highlighting the conditional “two-hit” nature of these phenotypes, and pointing to nucleotide imbalance as a potential biomarker of *LRRK2*-related PD progression. Furthermore, cell-type resolved analyses reveal that astrocytes and oligodendrocytes in the ventral midbrain are especially vulnerable to metabolic stress, displaying pronounced transcriptional downregulation of oxidative phosphorylation pathways in *G2019S LRRK2* mice. Together, these results emphasize the critical role of environmental factors, such as diet-induced metabolic syndrome, in unmasking latent vulnerabilities in PD-linked genetic backgrounds. Our data provide novel insight into potential metabolic targets for therapeutic intervention in individuals at-risk for developing PD.

## Methods Animals

*G2019S LRRK2* knockin mice (RRID: IMSR_JAX:030961), containing two base mutagenesis (GGG to AGC) in exon 41, were previously described [11]. The mice were purchased from Jackson Labs and maintained on a C57BL/6J background. Routine genotyping was done using a PCR-based method with primers (forward 5′-CAGGTAGGAGAACAAGTTTAC-3′, reverse 5′ -GGGAAAGCATTTAGTCTGAC-3′). The PCR produced a 307 bp band for the wild-type (WT) allele, a 383 bp band for the homozygous (HOMO) allele, and both bands for heterozygous (HET) samples. Male and female homozygous *G2019S LRRK2* knockin mice and their littermates were housed under specific pathogen-free conditions with constant temperature (22 °C), 30–70% humidity and a 12-hour light/dark cycle. Generally, mice were fed Purina Mills Lab Diet 5021 Autoclavable Breeder Mouse Diet and were provided with water *ad libitum*. Mice were routinely euthanized for experiments by i) cervical dislocation and decapitation (for obtaining fresh tissues) or by ii) lethal overdose with Avertin (2,2,2-tribromoethanol; i.p., 250 mg/kg) followed by cardiac perfusion with 4% paraformaldehyde solution (for obtaining fixed tissues). Mice were treated in accordance with the NIH Guidelines for the Care and Use of Laboratory Animals. All animal experiments were approved by the VAI Institutional Animal Care and Use Committee (IACUC).

### Experimental diet

Mice were fed with regular chow (Lab Diet 5021, 23.7% Kcal from fat) up to 11 months old, at which point they were randomly divided into either high-fat diet (D12492, 60% kcal fat, irradiated) or control diet (D12450J, 10% kcal fat, matching sucrose to D12492, irradiated) groups. The mice were maintained on their specific diet for an additional 5 months. Behavioral tests were conducted prior to the diet switch and again 4 months afterward. Body weight was measured biweekly. Fasting blood glucose and plasma insulin levels were measured 2 weeks prior to euthanasia. At the age of 16 months, plasma, brain, liver, lung, and kidney were collected for following analysis.

### Behavioral analysis

A dedicated mouse behavioral testing suite at Van Andel Institute was used for all behavioral assessments. ANY-maze video tracking system (version 7, Stoelting Co., https://www.any-maze.com/, RRID:SCR_014289) and CatWalk XT digital gait analysis system (version 10.6, Noldus, https://www.noldus.com/catwalk-xt, RRID:SCR_021262) were used to assess behavior during testing. Mice were habituated in the test room under normal lighting conditions for 1 hour prior to testing. A minimum rest period of 24 h was provided between assays. Behavioral apparatus used in testing was thoroughly cleaned with 70% ethanol between trials. For all assays, investigators were blinded to mouse genotype and treatment group.

#### Open field test

Prior to testing, mice were switched from normal lighting to red lighting. Mice were initially placed in a corner of the open field arena (50 x 50 x 40 cm, white) and monitored via an overhead placed video camera for 15 min. The center zone was defined as a 25 x 25 cm square in the middle of the box. Movement tracked by the camera was transmitted to a computer, where the ANY-maze software performed automated processing and analysis. Details of the open field test can be found at https://doi.org/10.1007/978-1-60761-303-9_1.

#### Elevated plus maze

Under normal lighting conditions, mice were initially placed in an enclosed corner of the maze apparatus (50 cm above the floor) and monitored via an overhead placed video camera for 10 min. Movement tracked by the camera was processed and analyzed by ANY-maze software. The entry into the open or closed arms was defined as full body entry based on body detection. Details of the Elevated Plus maze can be found at [43].

#### Y***-***maze

Under normal lighting conditions, mice were initially placed in the center of the maze apparatus and monitored with an overhead placed video camera for 10 min. Movement tracked by the camera was processed and analyzed by ANY-maze software. The entry into each of the three corridors of the Y-maze was defined as full body entry based on body detection. Percent alternation is defined as the non-repeat corridor entries relative to total entries. Details of the Y - maze can be found at https://www.any-maze.com/support/guides/automatically-scoring-spontaneous-alternations-in-the-y-maze-test/.

#### Catwalk digital gait analysis

Prior to testing, mice were switched from normal lighting to red lighting. Mice were initially placed on the walkway (40 cm), and 4 compliant runs were acquired of each animal and analyzed by CatWalk XT gait analysis system. Details of the Catwalk Gait Analysis can be found at https://noldus.com/catwalk-xt/resources.

### Blood analysis

Mice were fasted for 4 h before measurement, and the tail was cut at the tip and blood glucose levels were measured using a glucose meter (Cat# B07Z9N65F8, Accu-Chek). 20 μl blood was collected via capillary (Cat# 51684, The Lab Depot) from the tail vein and transferred into tubes containing EDTA as an anticoagulant (Sarstedt). Refer to protocol: https://doi.org/10.17504/protocols.io.81wgboy2ylpk/v1. The level of insulin in plasma was measured using the Ultra-Sensitive Mouse Insulin ELISA Kit (Cat# 90080, Crystal Chem) as per the manufacturer’s instructions.

### Tissue collection and processing

For histological analysis, mice were deeply anesthetized followed by transcardial perfusion with 0.9% NaCl and then 4% paraformaldehyde (PFA) in 0.1 M phosphate buffer (pH 7.4). Brains were collected and subsequently fixed for another 24 h in 4% PFA at 4 °C, followed by cryopreservation in 30% sucrose in 0.1 M PB for ≥ 24 h. Whole brains were subsequently processed using a microtome (Leica) into 35 μm-thick coronal sections, and serial sections of the whole brain were collected and used for subsequent staining. Refer to protocol: https://doi.org/10.17504/protocols.io.5jyl8pzk9g2w/v1. Peripheral tissues were collected and subsequently fixed for another 24 h in 4% PFA in 0.1 M phosphate buffer (pH 7.4), and fixed tissues were processed using automated tissue processors (Leica) and embedded in paraffin (Leica). Liver and lung were subsequently processed by a microtome (Leica) into 5 μm-thick sections.

For biochemical analysis, mice were anesthetized with an isoflurane vaporizer and whole blood was collected from cardiac puncture into a tube containing 10 µl of 0.5 M EDTA. Liver, lung, kidney and brain were then collected. Tissues were immediately snap frozen in liquid nitrogen and stored at –80 °C for later processing. Whole blood with EDTA was kept on wet ice until processing (<60 min) and centrifuged at 4,000 × g for 10 min at 4 °C. 50 μl plasma from each mouse was split and stored at -80 °C until analysis. Refer to protocol: https://doi.org/10.17504/protocols.io.eq2ly43kwlx9/v1.

### Immunohistochemistry and Hematoxylin-Eosin staining

Every 4^th^ serial brain section from each animal was pooled into a well for TH immunostaining (N300-109; Novus Biologicals), while every 8^th^ serial section was pooled for Iba1 (019-19741; FUJIFILM Wako) and GFAP (G3893; MilliporeSigma) immunostaining. FFPE lung sections were immunostained with CD68 (MCA1957GA, Biorad) and CD3 (ab11089, Abcam). Refer to protocols: https://doi.org/10.17504/protocols.io.3byl497zjgo5/v1, https://doi.org/10.17504/protocols.io.kxygx442dl8j/v1. Briefly, sections were quenched by incubation in 3% H_2_O_2_ (Sigma) diluted in methanol for 10 min at 4 °C, then blocked with 10% normal goat serum (Invitrogen) and 0.1% Triton-X100 in PBS for 1 h at room temperature. Sections were incubated with primary antibodies overnight at 4°C followed by incubation with biotinylated goat anti-mouse (BP-9200, Vector Laboratories) or goat anti-rabbit (BA-1000, Vector Laboratories) secondary antibodies for 2 h at room temperature. Sections were incubated with ABC reagent (PK-6100, Vector Laboratories) for 1 h at room temperature, color was developed by incubation with 3,3′-diaminobenzidine tetrahydrochloride (SK-4100, Vector Laboratories), and sections were mounted on Superfrost plus slides (#1255015, Fisher Scientific). Midbrain sections immunostained for TH were counterstained by incubating in cresyl violet solution for 15 min. Refer to **Table S7** for antibody details and dilutions used.

For Hematoxylin-Eosin (H&E) staining, slide-mounted liver and lung sections were processed using automated slide stainers (Leica) with optimized reagents (Leica).

### Digital slide scanning and analysis

Slide-mounted sections were scanned using an Aperio AT2 scanner (Aperio) at 20X magnification at a resolution of 0.5 μm/pixel. Annotation of hypothalamus, striatum and substantia nigra were performed using HALO analysis software (Indica Labs Inc., RRID:SCR_018350). GFAP-positive area, Iba1-positive microglial number and soma size were measured in the hypothalamus, striatum and substantia nigra using HALO analysis software (Area quantification and microglia activation modules; Indica Labs Inc., RRID:SCR_018350). TH-positive area in the striatum was measured using HALO analysis software (Area quantification module, Indica Labs Inc., RRID:SCR_018350). Lipid-positive area in the liver was measured using HALO analysis software (Area quantification module, Indica Labs Inc., RRID:SCR_018350).

### Stereological quantification of substantia nigra TH-positive neurons

Six midbrain sections (every 4^th^) of each animal were subjected to analysis. TH-positive and Nissl-positive neurons were counted using the optical fractionator probe of the StereoInvestigator software (MicroBrightField Biosciences, RRID:SCR_018948). Analysis area covered the entire substantia nigra pars compacta (SNpc). Random, systematic sampling was performed using a grid of 120 × 120 μm squares and applying an optical dissector with the dimensions 50 × 50 × 14 μm. Investigators were blinded to experimental conditions.

### Metabolite and lipid extractions

Frozen tissues were cryopulverized, and 30-40 mg aliquots from each tissue were split into pre-chilled 2 ml bead mill homogenizer tubes (19-627, Omni) and stored at -80 °C until analysis. All samples were homogenized using a ratio of 40 mg tissue per ml of solvent (1:1 chloroform:methanol) and processed using the Bligh-Dyer extraction method [44]. The same procedure was applied for plasma processing. The lower nonpolar phase of the resulting mixture was subjected to lipidomic analysis. The upper polar phase was subjected to metabolomic analysis. The RNA-containing interphase was processed for RNA isolation and subsequent RNA sequencing. Details of metabolite extraction can be found at [45].

### Lipidomic analysis

For lipidomics analysis, 14.5 µl plasma, 11.5 mg striatum, and 14.5 mg liver, kidney, and lung equivalents were aliquoted from the bottom nonpolar layer, dried under vacuum, and resuspended in 200 µl of ice-cold 1:1 acetonitrile (A955, Fisher):isopropanol (A461, Fisher). Deuterated PE 14:0 (14:0 PE-d54, 860371P, Avanti Research) was added as an internal standard to all lipidomics samples at a final concentration of 3 µg/ml to monitor instrument performance across each injection. Samples were analyzed with a Vanquish liquid chromatography system coupled with an Orbitrap ID-X (Thermo Fisher Scientific) using an H-ESI (heated electrospray ionization) source in positive or negative mode. Full scan data were collected using the Orbitrap with a scan range of 200-1700 m/z at a resolution of 240,000 and RF lens at 45%. Data-dependent MS2 fragmentation was induced in the Orbitrap using assisted higher-energy collisional dissociation (HCD) collision energies at 15, 30, 45, 75, and 110% as well as with collision-induced dissociation (CID) at a collision energy of 35%. For both MS2 fragmentations, Orbitrap resolution was 15,000 and the isolation window was 1.5 m/z. A m/z 184 mass trigger, indicative of phosphatidylcholines, was used for CID fragmentation. Data dependent MS3 fragmentation was induced in the ion trap with scan rate set at Rapid using CID at a collision energy of 35%. MS3 scans were triggered by specific acyl chain losses for detailed analysis of mono-, di-, and triacylglycerides. Total cycle time was 1.5 sec. LipidSearch software (version 5.0, Thermo Fisher, RRID:SCR_023716, https://www.thermofisher.com/us/en/home/industrial/mass-spectrometry/liquid-chromatography-mass-spectrometry-lc-ms/lc-ms-software/multi-omics-data-analysis/lipid-search-software.html) or Compound Discoverer software (version 3.3, Thermo Fisher) were used for initial lipid identification, and following analysis was conducted in Skyline (version 25.1, RRID:SCR_014080, https://skyline.ms/project/home/begin.view?). Detailed analysis process can be found at [45].

### Targeted metabolomic analysis

For metabolomics analyses, tissue samples underwent a second centrifugation step to ensure removal of tissue particles from the liquid metabolite layer. Finally, 32.8 µl plasma, 25.5 mg striatum, 26.1 mg liver, 32.4 mg kidney, and 30.2 mg lung equivalents were aliquoted from the second centrifugation of the top liquid layer, dried under vacuum, and resuspended in 200 µl or 100 µl of LC/MS-grade water for tissue and plasma samples, respectively. Deuterated glutamate (2,3,3,4,4-D₅, 98%, DLM-556, Cambridge Isotope Laboratories) internal standard was spiked in at resuspension at 10 μg/ml. Samples were analyzed with a Vanquish liquid chromatography system coupled to an Orbitrap Exploris 240 (Thermo Fisher Scientific) using an H-ESI source in negative mode. Full scan data were collected using the Orbitrap with a scan range of 70-850 m/z at a resolution of 240,000 and RF lens at 35%. Fragmentation was induced in the Orbitrap using assisted higher-energy collisional dissociation (HCD) collision energies at 15, 30, and 45%. Orbitrap resolution was 15,000, the isolation window was 2 m/z, and data-dependent scans were capped at 5 scans. Targeted mass MS2 triggers were included for a panel of compounds, displayed in **Table S8**. Peak picking and integration were conducted in Skyline (version 25.1). Detailed analysis process can be found at [45].

### Nucleoside quantitation

Absolute nucleoside quantitation was accomplished by running an external calibration curve with the brain and plasma samples. The stock mix contained Deoxyadenosine (dA) (27315, Caymen Chemical), Deoxycytidine (dC) (D3897, Sigma), Deoxyguanosine (dG) (9002864, Caymen Chemical), Deoxythymidine (dT) (20519, Caymen Chemical), and Deoxyuridine (dU) (D5412, Sigma) at a starting concentration of either 714 ng/ml (dC, dT, and dU) or 71.4 ng/ml (dG and dA), followed by an 8 step 2x serial dilution for nine total curve points. An isotopically labeled standard, ^13^C_10_-^15^N_2_-2’Dexoythymidine (CNLM-3902, Cambridge Isotope Laboratories), was spiked in at resuspension at a concentration of 20 ng/ml. Samples were analyzed with an Agilent 6470 triple quadrupole mass spectrometer coupled with an Agilent ultra-high performance liquid chromatography 1290 Infinity II. Refer to supplementary methods for detailed parameters. Peak picking was performed using Agilent Masshunter Quantitative Analysis software (Version B.09.00/Build 9.0.647.0 for QQQ, RRID:SCR_015040, http://www.agilent.com/en-us/products/software-informatics/masshunter-suite/masshunter/masshunter-software). The peak areas from the curves were used to generate linear regression for quantitation of the target analytes.

### RNA extraction and bulk sequencing

RNA was isolated from brain tissue (following metabolite extraction [46]) and pulverized liver tissue using RNeasy Plus kits (Qiagen) per manufacturer’s instruction, and total RNA libraries were prepared from 500 ng RNA using Kapa RNA HyperPrep (Roche) with Qiaseq Fastselect (Qiagen). RNA libraries were normalized, pooled and sequenced on Illumina NovaSeq 6000 using paired end, 50 base pair reads to an average depth of 50M reads per sample. Sequencing data were analyzed following the bulk RNAseq workflow at https://doi.org/10.5281/zenodo.15651884. Details of RNA sequencing can be found at https://www.illumina.com/techniques/sequencing/rna-sequencing.html.

### Single-nuclei RNA sequencing and data analysis

Single nuclei were isolated from the mouse ventral midbrain using the protocol described at https://doi.org/10.17504/protocols.io.14egnr8kml5d/v1. The purified nuclei were then fixed using the Chromium Next GEM Single Cell Fixed RNA Sample Preparation Kit (1000414, 10x Genomics), following the manufacturer’s instructions. Library preparation was performed using the Chromium Fixed RNA Kit (1000496, 10x Genomics), also according to the manufacturer’s protocol. 16 libraries were pooled and sequenced on Illumina NovaSeq 6000 using paired end, 100 base pair read to an average depth of 80M reads per sample.

Fastq files were processed using the ‘cellranger multi’ workflow in 10x CellRanger (v7.1.0 , RRID:SCR_017344, https://support.10xgenomics.com/single-cell-gene-expression/software/pipelines/latest/what-is-cell-ranger) with ‘refdata-gex-mm10-2020-A’ reference and the ‘v1.0.1_mm10-2020-A’ probe set. The filtered gene counts were processed using a standard scRNA-seq workflow as implemented in the R (v4, RRID:SCR_001905, https://www.r-project.org/) package, scater (v1.26.1, RRID:SCR_015954, https://bioconductor.org/packages/release/bioc/html/scater.html), and its associated packages including scran (v1.26.2, RRID:SCR_016944, https://bioconductor.org/packages/release/bioc/html/scran.html), scuttle (v1.8.4, https://bioconductor.org/packages/release/bioc/html/scuttle.html) and batchelor (v1.14.1, https://www.bioconductor.org/packages/release/bioc/html/batchelor.html) [47–49]. Specifically, nuclei were filtered based on low library size, low number of genes detected and high mitochondrial gene count percentage. The filter thresholds were calculated for each sample separately as five median-absolute-deviations (MAD); for the high mitochondrial count percentage threshold, the cutoff was required to be at least 0.5%.

The deconvolution approach for normalization was performed using the quickCluster, computeSumFactors and logNormCounts functions with default parameters. The quickCorrect wrapper function was run with ‘batch’ set to each sample and ‘PARAM=FastMnnParam(auto.merge=TRUE)’; this normalizes the gene counts, identifies the top most variable genes and runs mutual nearest neighbor (MNN) correction using the FastMNN algorithm [50]. Clusters were identified by constructing a k-means for k-nearest neighbors graph based on the MNN output and running the walktrap clustering algorithm. The corrected reduced dimensions were also used for tSNE as implemented in the runTSNE function.

Doublet scoring was done for each sample separately since doublets are formed between cells/nuclei of the same library. Each sample was reprocessed from normalization to clustering as described above except no batch correction was done and the first 50 principal components, from a PCA of the top 2000 most variable genes, were used for clustering. Two doublet scoring algorithms, scDblFinder (RRID:SCR_022700, https://github.com/plger/scDblFinder) and computeDoubletDensity, from the scDblFinder package v1.12.0 [51] were run to assign doublet scores to each nucleus, which were appended to the combined analysis above. Visual inspection of doublet scores from both methods identified a large doublet cluster containing 14004 nuclei out of a total of 48700. This doublet cluster was removed and the normalization, batch correction, dimension reductions and clustering were rerun as described above.

Automated cell type annotations were performed using the SingleR (RRID:SCR_023120, https://www.bioconductor.org/packages/release/bioc/html/SingleR.html) function from the SingleR package v2.0.0, with “de.method = ‘wilcox’” [52]. The reference dataset was midbrain nuclei from the downsampled Mouse Brain Atlas dataset hosted on https://singlecell.broadinstitute.org/ as accession SCP2158 [53]. Log normalized counts were used for both the query and the reference datasets.

### Western blot analysis

Brain, liver tissue and lung tissues were homogenized using a tissue homogenizer at maximum speed for 10 sec in chilled lysis buffer (50 mM Tris-HCl Ph7.5, 1 mM EGTA, 1% Triton-X 100, 0.27 M sucrose, phosphatase inhibitor cocktail set 1 (#524624, EMD Millipore), phosphatase inhibitor cocktail 3 (#P0044, Sigma-Aldrich), protease inhibitor cocktail (#11836170001, Sigma-Aldrich)), followed by incubation on ice for 15 min. The samples were then centrifuged at 4°C for 30 min at 21,000 rcf. Supernatants were saved for following analysis. Protein concentrations were measured using a Pierce BCA protein assay following the manufacturer’s instructions (Thermofisher Scientific). 80 μg protein of each sample was loaded into SDS-PAGE gels and run at 120V, followed by transferring to nitrocellulose membranes (0.2 µm) at 25 V overnight. For detailed protocol, see https://doi.org/10.17504/protocols.io.261ge545jg47/v1. Primary antibodies used for Western blotting include: LRRK2 (75-253, NeuroMab), pS935-LRRK2 (ab133450, Abcam), pS1292-LRRK2 (ab203181, Abcam), Rab8 (D22D8, Cell Signaling Technology), pT72-Rab8a (ab230260, Abcam), Rab10 (8127S; Cell Signaling Technology), pThr73-Rab10 (ab230261, Abcam), Rab12 (18843-1-AP, Protein Tech), pSer106-Rab12 (ab256487, Abcam), Actin (MAB1501, Millipore), Thymidine Phosphorylase (AF7568, Novus Biologicals). Refer to **Table S7** for antibody details and dilutions used. Images were acquired using a ChemiDoc System (Bio-Rad) and analyzed using Bio-Rad Image Lab Software (RRID:SCR_014210).

### Proteomic analysis

The pulverized lung tissues were homogenized using a Bead Ruptor Elite (Cat# 19-042E, Omni International) for 30 sec in 4% SDS solution containing 1x HALT Protease (Cat# 78442, Thermo Fisher Scientific). Samples were sonicated and clarified via centrifugation and transferred to a Protein LoBind Eppendorf tube. Proteins were quantified using the Pierce BCA Protein Assay kit (Cat# 23227, Thermo Fisher Scientific) and 100 µg of protein was aliquoted for digestion. Protein digestion utilized the S-Trap (Cat# CO2-Mini, Protifi) platform to remove any SDS prior to LC-MS/MS analysis. Briefly, proteins were reduced with dithiothreitol (DTT) for 20 min, alkylated with iodoacetamide (IAA) for 20 min and digested overnight with Trypsin/Lys-C (Cat# V5072, Promega) at a ratio of 50:1 (protein:enzyme (w/w)). Peptides were eluted off the S-Trap Column and dried in a Genevac SpeedVac. Dried samples were then cleaned up with C-18 reverse phase columns from Harvard Apparatus (Cat# 74-4601) and dried before resuspension for LC-MS/MS analysis. Dried samples were resuspended in 50 µl 0.1% FA (LS118-1, Fisher Scientific) and diluted with 50 µl of 0.1% TFA (LS119-500, Fisher Scientific).

Data-independent Acquisition (DIA) analyses were performed on an Orbitrap Eclipse coupled to Vanquish Neo system (Thermo Fisher Scientific) with a FAIMS Pro source (Thermo Fisher Scientific) located between the nanoESI source and the mass spectrometer. Full MS spectra were collected at 120,000 resolution (full width half-maximum; FWHM), and MS2 spectra at 30,000 resolution (FMWH). Both the standard automatic gain control (AGC) target and the automatic maximum injection time were selected. A precursor range of 380-985 m/z was set for MS2 scans, and an isolation window of 50 m/z was chosen with a 1 m/z overlap for each scan cycle. 32% HCD collision energy was used for MS2 fragmentation. DIA data was processed in Spectronaut (version 18, Biognosys, Switzerland, https://biognosys.com/software/spectronaut/) using direct DIA. Data was searched against the *Mus musculus* proteome, including expected mutations. The manufacturer’s default parameters were used. Briefly, trypsin/P was set as digestion enzyme and two missed cleavages were allowed. Cysteine carbamidomethylation was set as fixed modification, and methionine oxidation and protein N-terminus acetylation as variable modifications. Identification was performed using a 1% q-value cutoff on precursor and protein levels. Both peptide precursors and protein false discovery rate (FDR) were controlled at 1%. Ion chromatograms of fragment ions were used for quantification. For each targeted ion, the area under the curve between the XIC peak boundaries was calculated. Detailed data analysis process can be found at [45].

### cDNA synthesis and qPCR

RNA was isolated from mouse ventral midbrain tissue using the RNeasy Plus kit (#74104, Qiagen) per manufacturer’s instruction. RNA concentration and purity were determined using a spectrophotometer (NanoDrop, Thermo Scientific). cDNA was synthesized from 1 μg total RNA per sample using SuperScriptTM III Reverse Transcriptase (#18080, Invitrogen) per manufacturer’s instruction. qPCR was carried out on a real-time PCR system (QuantStudio 3, Thermo Fisher Scientific) using SYBR Green detection chemistry. 100 ng of cDNA was used as the template. Primers targeting TYMP (For - CCGAGCCAGATCCAGAAAACTG -, Rev - GCTTCTCTCAGATGCCCTCCAT -) were used to assess transcript abundance. ACTB (β-actin, For - GTACTCTGTGTGGATCGGTGG -, Rev – GCAGCTCAGTAACAGTCCG -) served as the endogenous reference gene. Relative gene expression was calculated using the ΔΔCt method, with ACTB providing normalization. For detailed protocol, see http://doi.org/10.17504/protocols.io.bp2l6zjwrgqe/v1.

### Statistical analysis

Data from behavior test, histological analysis, Western blot analysis and selected metabolomics experiments was analyzed with GraphPad Prism 10 (RRID:SCR_002798) software by unpaired Student’s *t*-test or two-way ANOVA with Tukey’s multiple comparisons test. Graphs were generated using GraphPad Prism 10 and depict all data as mean ± SEM.

Differential abundance analyses of metabolites and lipids were performed using R v4.4.0 using workflows adapted from methods published here [46]. For each tissue and compound type (i.e. metabolite/lipid), limma-eBayes was fit on log_2_ transformed abundances. The design matrix included both genotypes, both diets, and their interaction. Compounds were imputed using QRLIC, and compounds with >30% of values missing were removed prior to analysis. Benjamini-Hochberg multiple testing corrections were used to control the false discovery rate at 5%. Process, analysis, and visualization code is available at https://zenodo.org/records/18166240.

## Supporting information

Knowledge Resource Table

## List of abbreviations

PD: Parkinson’s disease
MetS: Metabolic syndrome
HFD: High-fat diet
Con.diet: Control diet
TP: Thymidine phosphorylase
H&E: Hematoxylin-Eosin
PPP: Pentose Phosphate Pathway
TCA: citric acid cycle
2-KG: 2-Ketoglutarate

## Availability of data, code and materials

The data, code, protocols, and key lab materials used and generated in this study are listed in a Key Resource Table alongside their persistent identifiers at https://doi.org/10.5281/zenodo.17860645. Behavioral test videos were deposited in the BioStudies database (https://www.ebi.ac.uk/biostudies/studies/S-BSST2573). Microscopy images were deposited in the Brain Image Library (https://doi.org/10.35077/g.1191). Raw data generated by metabolomics, lipidomics, proteomics, bulk RNA-seq, and single-nucleus RNA-seq in this study have been deposited in the ASAP CRN Cloud. An earlier version of this manuscript was posted to *bioRxiv* on 2026-01-02 at https://doi.org/10.64898/2025.12.31.697225.

## Acknowledgements

We would like to thank the outstanding core facilities at the Van Andel Institute that supported this work, including the Vivarium Core (RRID:SCR_023211), Pathology and Biorepository Core (RRID:SCR_022912), Kaitlyn DenHaan from the Bioinformatics and Biostatistics Core (RRID:SCR_024762), Katelyn Becker from the Genomics Core (RRID:SCR_022913), and Colt Capan, Hyoungjoo Lee, Ryan Sheldon, Christine Isaguirre, and Abigail E. Ellis from the Mass Spectrometry Core (RRID:SCR_024903). Figure 1a, Figure 10a, and Figure 12 were generated using BioRender. This work was supported by the joint efforts of The Michael J. Fox Foundation for Parkinson’s Research (MJFF) and the Aligning Science Across Parkinson’s (ASAP) initiative. MJFF administers the grant (ASAP-000592, to DJM) on behalf of ASAP and itself. Yue Ma was supported by a Van Andel Institute - West Michigan Neurodegenerative Diseases (MiND) Program Pathway-to-Independence award (MiP2i.001.YM).

## Author contributions

YM designed, performed, and analyzed most experiments. MLE performed stereological analysis. KS, AO and JN assisted with sample collection and polarization. ZBM assisted with Mass Spec data analysis. ZF assisted with bulk RNA-seq data analysis. KHL assisted with snRNAseq data analysis. DJM conceptualized and supervised this research. YM and DJM wrote the manuscript.

