## Supplementary Data for "Interaction of *G2019S LRRK2* and metabolic syndrome in a two-hit mouse model of Parkinson’s disease: LRRK2-driven systemic depletion of pyrimidine nucleosides"

### Supplementary Figures

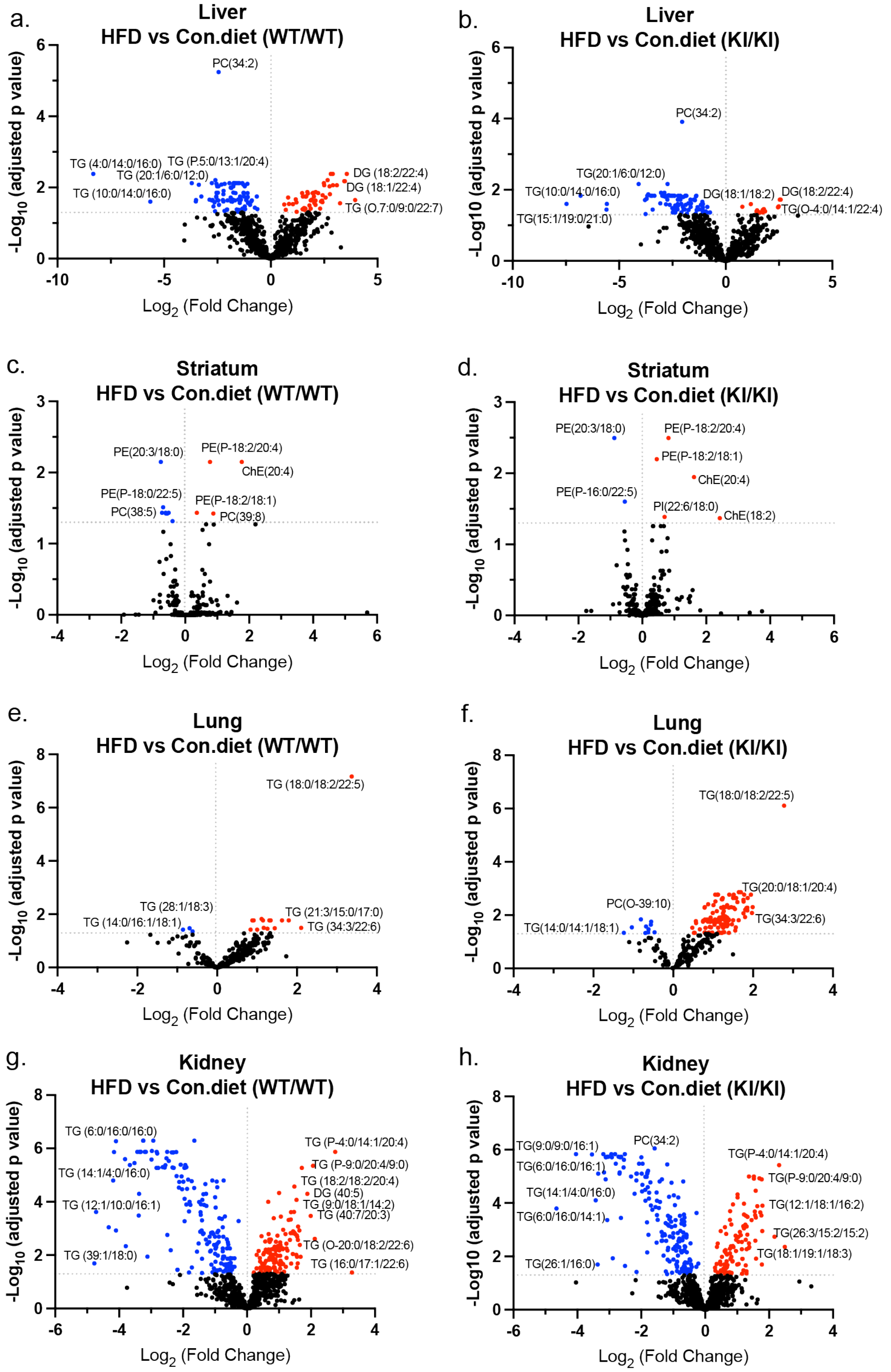

**Figure S1. Lipid alterations across tissues in *G2019S LRRK2* knockin and wild-type mice under HFD.** Volcano plot showing differences in the abundance of lipids between HFD and Con. diet in liver (**a-b**), striatum (**c-d**), lung (**e-f**), and kidney (**g-h**) of *G2019S LRRK2* knockin mice and wild-type controls. WT/WT, wild-type mice; KI/KI, homozygous *G2019S LRRK2* knockin mice. Each point represents a lipid species. X-axis shows $\log_2$ (fold change) in lipid abundance between high-fat diet and control diet groups. Y-axis shows $-\log_{10}$ (adjusted *p*-value) from limma's empirical Bayes moderated analysis (eBayes) combined with Benjamini-Hochberg (BH) for multiple testing correction ($n$ = 6 per group). Lipids with adjusted $P<0.05$ are considered significantly altered. Significantly upregulated lipids are highlighted in red and significantly downregulated lipids are highlighted in blue.

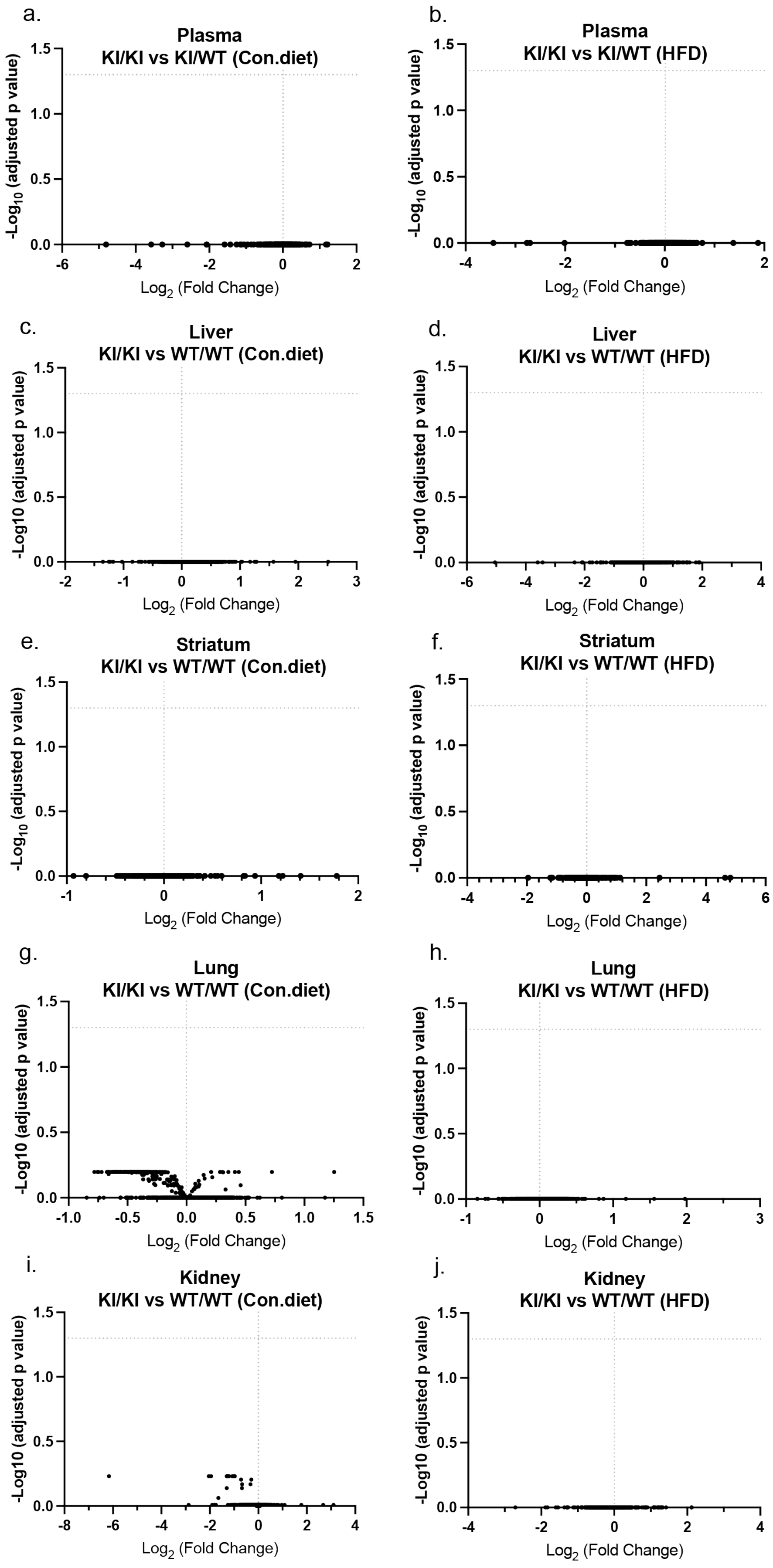
a.
Plasma
KI/KI vs KI/WT (Con.diet)
b.
Plasma
KI/KI vs KI/WT (HFD)
c.
Liver
KI/KI vs WT/WT (Con.diet)
d.
Liver
KI/KI vs WT/WT (HFD)
e.
Striatum
KI/KI vs WT/WT (Con.diet)
f.
Striatum
KI/KI vs WT/WT (HFD)
g.
Lung
KI/KI vs WT/WT (Con.diet)
h.
Lung
KI/KI vs WT/WT (HFD)
i.
Kidney
KI/KI vs WT/WT (Con.diet)
j.
Kidney
KI/KI vs WT/WT (HFD)
$-Log_{10}$ (adjusted p value)
-Log10 (adjusted p value)
$Log_2$ (Fold Change)

**Figure S2. Lipid alterations across tissues in *G2019S LRRK2* knockin mice under HFD and control diet.** Volcano plot shows differences in the abundance of lipids between *G2019S LRRK2* knockin and wild-type mice in the plasma (a, b), liver (c-d), striatum (**e-f**), lung (**g-h)**, and kidney (**i-j**) under HFD and Con. diet. WT/WT, wild-type mice; KI/KI, homozygous *G2019S LRRK2* knockin mice. Each point represents a lipid species. X-axis shows $\log_2$ (fold change) in lipid abundance between high-fat diet and control diet groups. Y-axis shows $-\log_{10}$ (adjusted *p*-value) from limma's empirical Bayes moderated analysis (eBayes) combined with Benjamini-Hochberg (BH) for multiple testing correction ($n$ = 6 per group). Lipids with adjusted $P<0.05$ are considered significantly altered. No significant alterations were found.

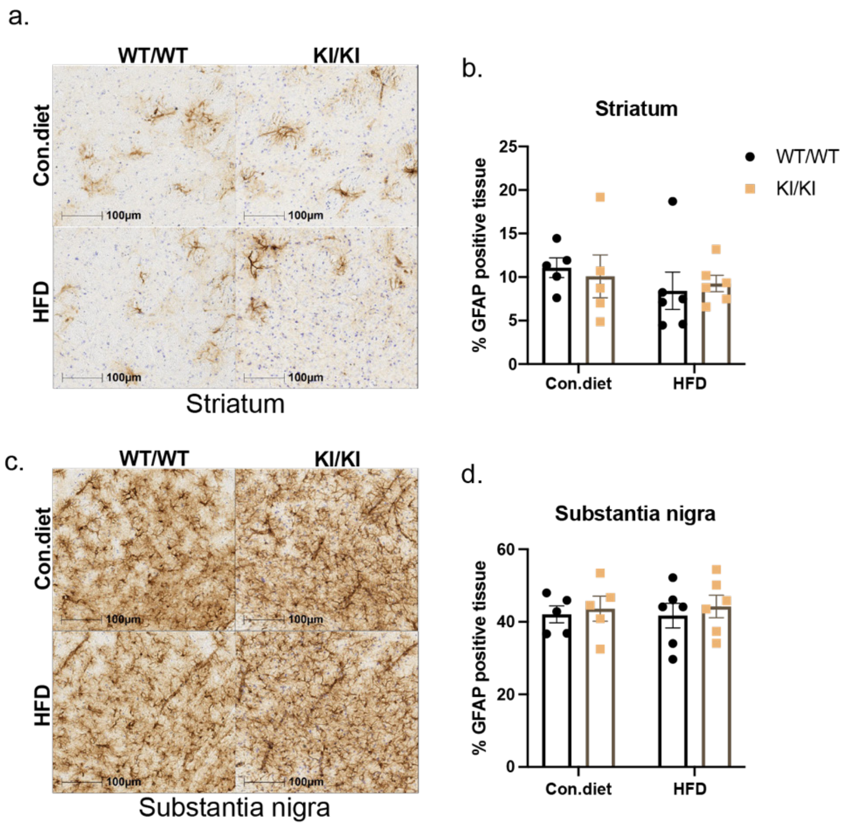

**Figure S3. Pre-existing metabolic syndrome fails to induce reactive astrogliosis in striatum and substantia nigra of *G2019S LRRK2* knockin mice. a, c** Representative image of GFAP immunostaining in the striatum (a) and substantia nigra (c) showing astrocytes distribution. Scale bar: 100 μm. **b, d** Quantification of GFAP-positive area in the striatum (b) and substantia nigra (d) (*n* = 5-6 per group). WT/WT, wild-type mice; KI/KI, homozygous *G2019S LRRK2* knockin mice. Data are presented as mean ± SEM. A two-way ANOVA was performed to evaluate the effects of high-fat diet and *G2019S LRRK2* and their interaction followed by Fisher's LSD comparisons. No significant effects were found.

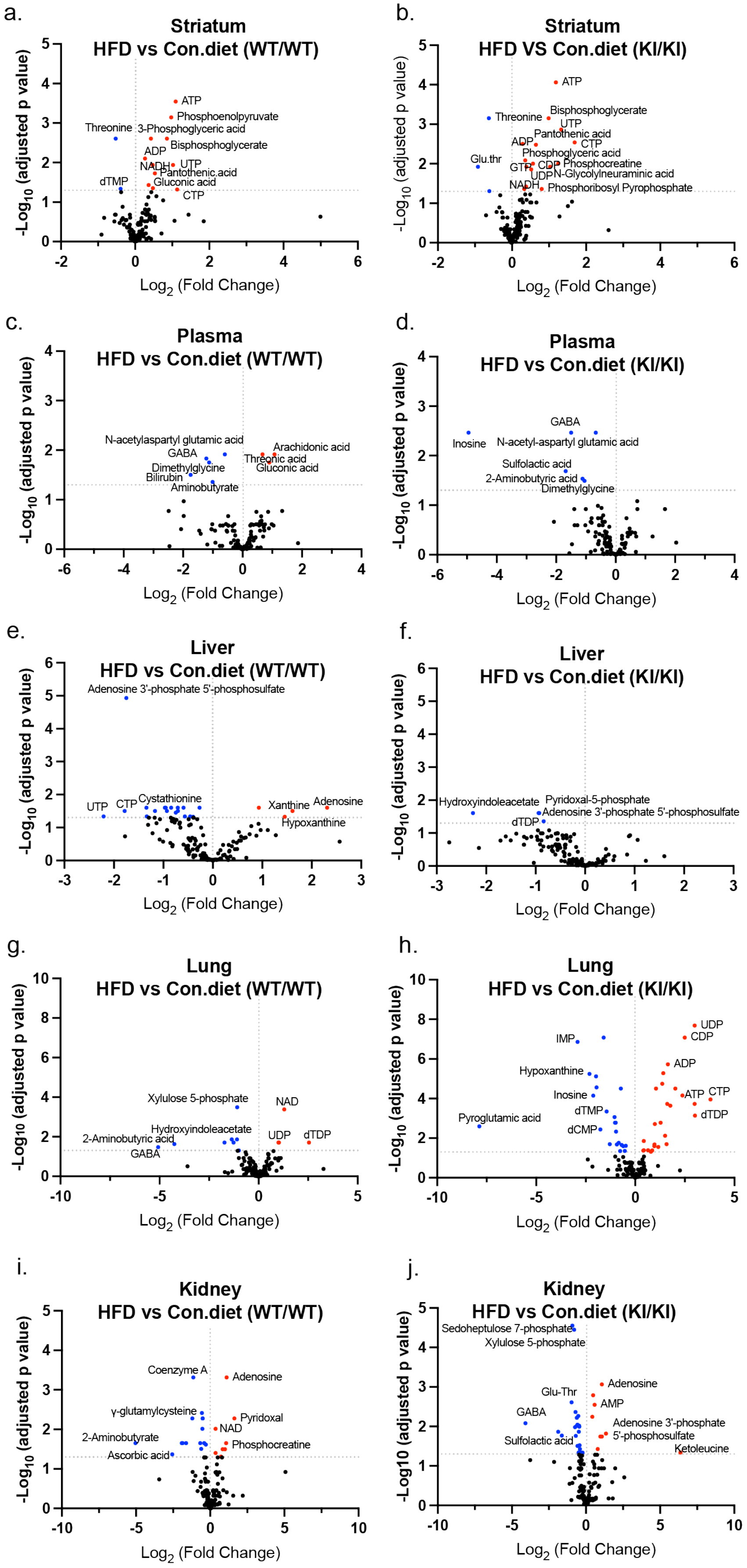
a.
Striatum
HFD vs Con.diet (WT/WT)
-Log10 (adjusted p value)
Log2 (Fold Change)
ATP
Phosphoenolpyruvate
Threonine
3-Phosphoglyceric acid
Bisphosphoglycerate
ADP
NADH
UTP
Pantothenic acid
dTMP
Gluconic acid
CTP
b.
Striatum
HFD VS Con.diet (KI/KI)
-Log10 (adjusted p value)
Log2 (Fold Change)
ATP
Bisphosphoglycerate
Threonine
UTP
Pantothenic acid
ADP
CTP
Phosphoglyceric acid
Glu.thr
GTP
CDP
Phosphocreatine
UDP
N-Glycolylneuraminic acid
NADH
Phosphoribosyl Pyrophosphate
c.
Plasma
HFD vs Con.diet (WT/WT)
-Log10 (adjusted p value)
Log2 (Fold Change)
N-acetylaspartyl glutamic acid
Arachidonic acid
GABA
Threonic acid
Dimethylglycine
Gluconic acid
Bilirubin
Aminobutyrate
d.
Plasma
HFD vs Con.diet (KI/KI)
-Log10 (adjusted p value)
Log2 (Fold Change)
GABA
Inosine
N-acetyl-aspartyl glutamic acid
Sulfolactic acid
2-Aminobutyric acid
Dimethylglycine
e.
Liver
HFD vs Con.diet (WT/WT)
-Log10 (adjusted p value)
Log2 (Fold Change)
Adenosine 3'-phosphate 5'-phosphosulfate
UTP
CTP
Cystathionine
Xanthine
Adenosine
Hypoxanthine
f.
Liver
HFD vs Con.diet (KI/KI)
-Log10 (adjusted p value)
Log2 (Fold Change)
Hydroxyindoleacetate
Pyridoxal-5-phosphate
Adenosine 3'-phosphate 5'-phosphosulfate
dTDP
g.
Lung
HFD vs Con.diet (WT/WT)
-Log10 (adjusted p value)
Log2 (Fold Change)
Xylulose 5-phosphate
NAD
Hydroxyindoleacetate
2-Aminobutyric acid
UDP
dTDP
GABA
h.
Lung
HFD vs Con.diet (KI/KI)
-Log10 (adjusted p value)
Log2 (Fold Change)
UDP
CDP
IMP
ADP
Hypoxanthine
Inosine
ATP
CTP
dTMP
Pyroglutamic acid
dTDP
dCMP
i.
Kidney
HFD vs Con.diet (WT/WT)
-Log10 (adjusted p value)
Log2 (Fold Change)
Coenzyme A
Adenosine
γ-glutamylcysteine
Pyridoxal
NAD
2-Aminobutyrate
Phosphocreatine
Ascorbic acid
j.
Kidney
HFD vs Con.diet (KI/KI)
-Log10 (adjusted p value)
Log2 (Fold Change)
Sedoheptulose 7-phosphate
Xylulose 5-phosphate
Adenosine
Glu-Thr
AMP
GABA
Adenosine 3'-phosphate
5'-phosphosulfate
Sulfolactic acid
Ketoleucine

**Figure S4. Non-lipid metabolite alterations across tissues in *G2019S LRRK2* knockin and wild-type mice under HFD.** Volcano plot shows differences in the abundance of non-lipid metabolites between HFD and Con. diet groups in *G2019S LRRK2* knockin mice and wild-type mice in the striatum (**a, b**), plasma (**c-d**), liver (**e-f**), lung (**g-h)**, and kidney (**i-j**). WT/WT, wild-type mice; KI/KI, homozygous *G2019S LRRK2* knockin mice. Each point represents a non-lipid metabolite. X-axis shows $\log_2$ (fold change) in non-lipid metabolite abundance between HFD and Con. diet groups. Y-axis shows $-\log_{10}$ (adjusted *p*-value) from limma's empirical Bayes moderated analysis (eBayes) combined with Benjamini-Hochberg (BH) for multiple testing correction ($n$ = 6 per group). Non-lipid metabolites with adjusted $P<0.05$ are considered significantly altered. Significantly upregulated metabolites are highlighted in red and significantly downregulated metabolites are highlighted in blue.

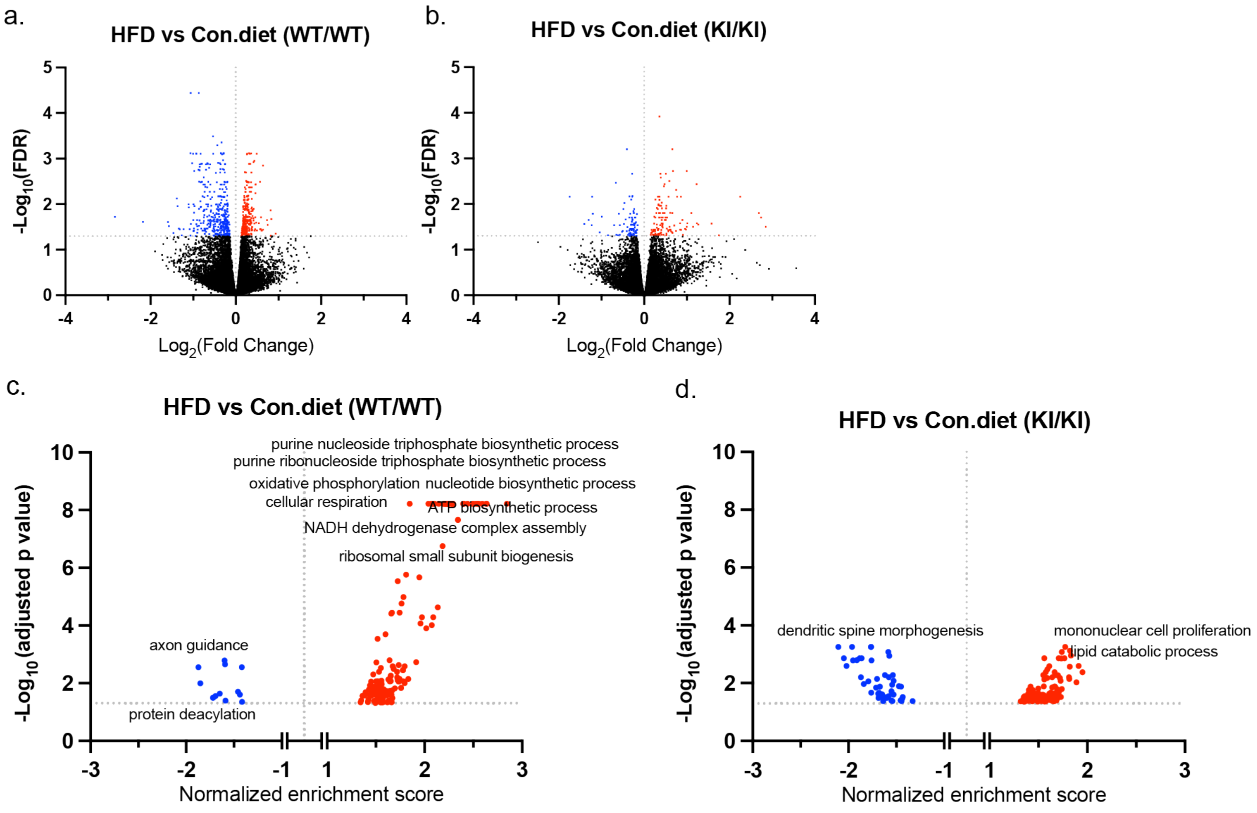

**Figure S5. Pre-existing metabolic syndrome reveals altered brain energy metabolism in *G2019S LRRK2* knockin mice. a-b** Volcano plot showing differential gene expression in the striatum between HFD and Con.diet in *G2019S LRRK2* knockin and wild-type mice. Each point represents one gene. X-axis shows $\log_2$ (fold change) in gene expression. Y-axis shows $-\log_{10}$ (adjusted *p*-value), significance was defined as a Benjamini–Hochberg adjusted $P<0.05$ ($n = 4$ per group). Gene expression with adjusted $P<0.05$ are considered significantly altered. Significantly upregulated genes are highlighted in red and significantly downregulated genes are highlighted in blue. **c-d** Volcano plot illustrates enriched biological processes between *G2019S LRRK2* knockin and wild-type mice under Con.diet and HFD conditions as identified by gene set enrichment analysis (GSEA). Each point represents one process, X-axis shows normalized enrichment score, Y-axis shows $-\log_{10}$ (adjusted *p*-value), significance was defined as a Benjamini–Hochberg adjusted $P<0.05$ ($n = 4$ per group). Significantly upregulated processes are highlighted in red and significantly downregulated processes are highlighted in blue. WT/WT, wild-type mice; KI/KI, homozygous *G2019S LRRK2* knockin mice.

a.

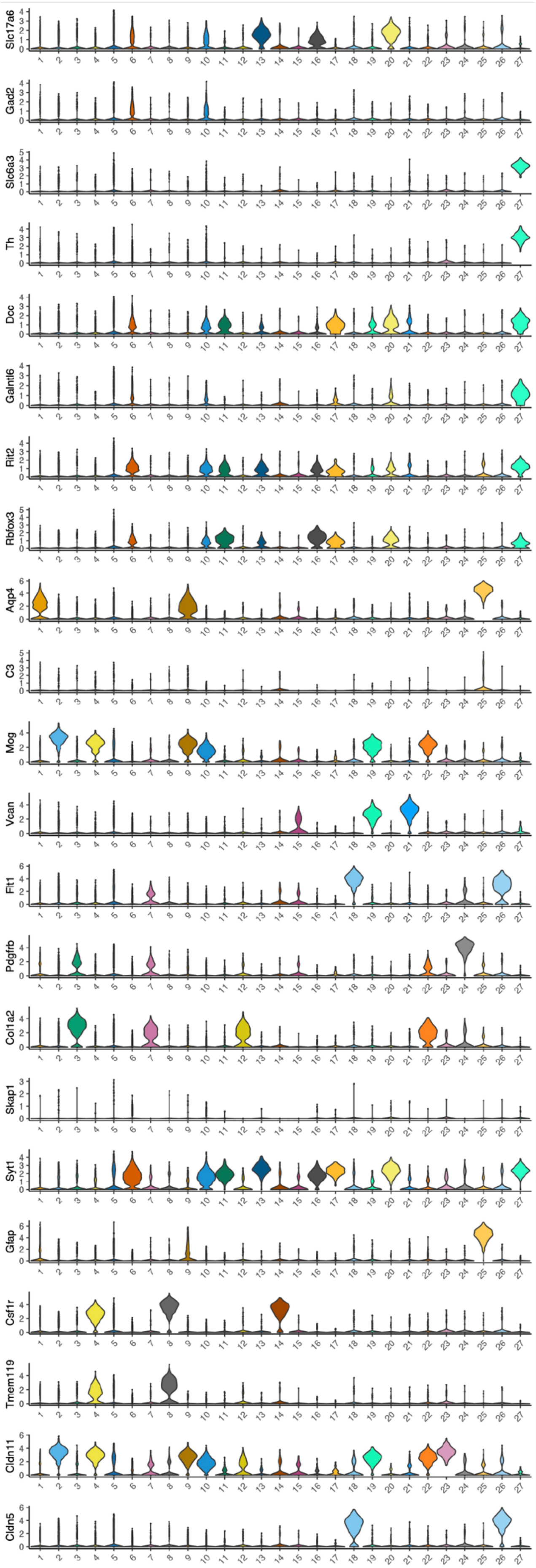

b.

| Cell type | Cluster | Counts | Sum |
|---|---|---|---|
| Oligo | 2 | 6301 | 12293 |
| Oligo | 23 | 96 | |
| Oligo hibrid | 22 | 472 | |
| Oligo hibrid | 4 | 1155 | |
| Oligo hibrid | 10 | 2820 | |
| Oligo hibrid | 9 | 1449 | |
| Neuron | 6 | 6192 | 8015 |
| Neuron | 11 | 217 | |
| Neuron | 13 | 242 | |
| Neuron | 16 | 577 | |
| Neuron | 17 | 115 | |
| Neuron | 20 | 534 | |
| Neuron | 27 | 138 | |
| Astrocyte | 1 | 3880 | 4294 |
| Asttrocyte | 25 | 414 | |
| Fibroblast | 3 | 1460 | 2216 |
| Fibroblast | 7 | 318 | |
| Fibroblast | 12 | 438 | |
| OPC | 15 | 149 | 1653 |
| OPC | 19 | 387 | |
| OPC | 21 | 1117 | |
| Microglia | 8 | 1115 | 1297 |
| Microglia | 14 | 182 | |
| Endothelia | 18 | 537 | 785 |
| Endothelia | 26 | 248 | |
| Pericyte | 24 | 209 | 209 |
| no annotate | 5 | 3934 | 3934 |

**Figure S6. Identification of cell type-specific clusters and numbers in the mouse ventral midbrain via snRNA-seq analysis.** Nuclei were isolated from ventral midbrain of *G2019S LRRK2* knockin mice and their WT littermates (*n* = 4 mice per group), with 34,969 nuclei recovered in total. **a** Cell type annotation of each cluster using well established cell-type transcript markers. **b** Cell counts and cell type annotated in each cluster.

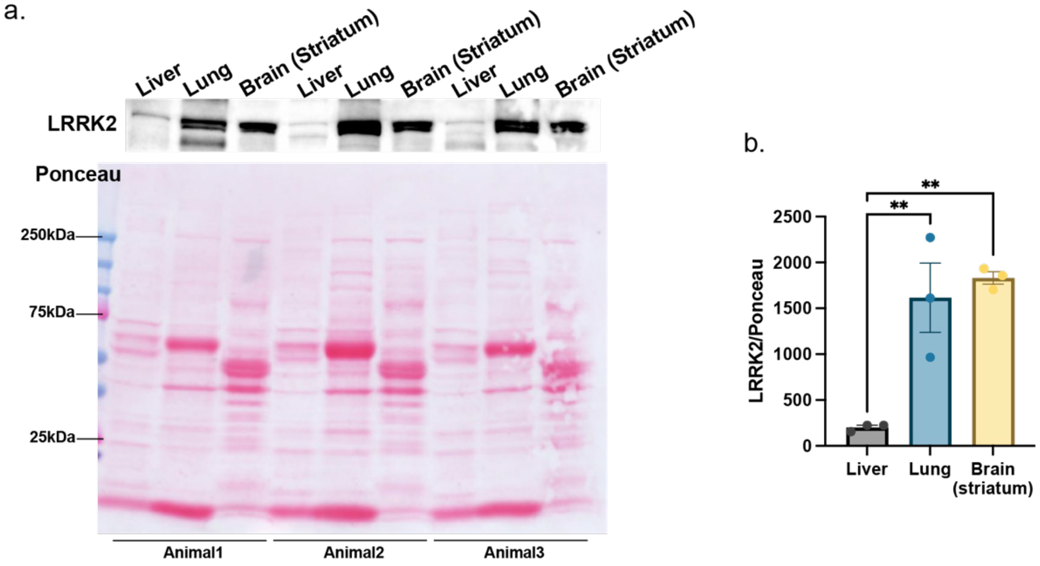

**Figure S7. Western blot analysis and quantification of LRRK2 protein levels across tissues. a** Sixty µg of total protein from liver, lung, and striatum of adult wild-type mice (*n* = 3 mice/group) were resolved by SDS-PAGE and transferred to nitrocellulose. Membrane was stained with Ponceau S to assess total protein loading, followed by immunoblotting with an anti-LRRK2 antibody. **b** Quantification of LRRK2 protein levels normalized to Ponceau signal. Data are presented as mean ± SEM (*n* = 3 mice/group). One-way ANOVA was performed to evaluate the difference among groups. Significant differences were observed (***P*<0.01) relative to liver.

### Supplementary Tables

**Table S1**. Summary of metabolites and lipids identified by metabolomic and lipidomic analysis.

**Table S2**. Differentially-expressed genes identified by bulk RNA-seq analysis of the striatum.

**Table S3**. Enriched GO Biological Process (GOBP) terms identified from bulk RNA-seq analysis of the striatum.

**Table S4**. Differentially-expressed genes identified by single-nucleus RNA-seq analysis of the ventral midbrain.

**Table S5**. Enriched GO Biological Process (GOBP) terms identified from single-nucleus RNA-seq analysis of the ventral midbrain.

**Table S6.** Summary of relative protein abundance quantified by mass spectrometry in lung tissue.

**Table S7.** Antibody information for Western blot (WB) and immunohistochemistry (IHC).

**Table S8.** Panel of compounds targeted for MS2 fragmentation.

**Key Resource Table (KTR).**

**Table S7.** Antibody information for Western blot (WB) and immunohistochemistry (IHC).

| **Target (WB)** | **Catalog no.** | **Host** | **Dilution** |
|---|---|---|---|
| LRRK2 (N241A/34) | 75-253, NeuroMab | mouse | 1:1000 |
| pSer935-LRRK2 | ab133450, Abcam | rabbit | 1:1000 |
| pSer1292-LRRK2 | ab203181, Abcam | rabbit | 1:1000 |
| Rab8a | D22D8, Cell Sigal Tech | rabbit | 1:1000 |
| pThr72-Rab8a | ab230260, Abcam | rabbit | 1:500 |
| Rab10 | 8127S, Cell Signal Tech | rabbit | 1:1000 |
| pThr73-Rab10 | ab230261, Abcam | rabbit | 1:500 |
| Rab12 | 18843-1-AP, Proteintech | rabbit | 1:1000 |
| pSer106-Rab12 | ab256487, Abcam | rabbit | 1:1000 |
| Thymidine Phosphorylase | AF7568, R&D systems | sheep | 1:200 |
| Actin | MAB1501, Millipore | mouse | 1:2000 |
| **Target (IHC)** | **Catalog no.** | **Host** | **Dilution** |
| Tyrosine Hydroxylase (TH) | N300-109, Novus Biologicals | rabbit | 1:2000 |
| Iba1 | 019-19741, FUJIFILM Wako | rabbit | 1:1000 |
| GFAP | G3893, Millipore Sigma | mouse | 1:1000 |
| CD68 | MCA1957GA, Biorad | rat | 1:500 |
| CD3 | ab11089, Abcam | rat | 1:200 |

**Table S8.** Panel of compounds targeted for MS2 fragmentation.

| Compound | Formula | Adduct | m/z | z | RT Time (min) | Window (min) |
|---|---|---|---|---|---|---|
| **2,3-Dihydroxy-2-methylbutanoic acid** | C5H10O4 | -H | 133.0506 | 1 | 8.78 | 4 |
| **D-2-Aminobutyrate** | C4H9NO2 | -H | 102.0561 | 1 | 1.2 | 2 |
| **Deoxyuridine** | C9H12N2O5 | -H | 227.0673 | 1 | 2.1 | 4 |
| **2-Hydroxybutyrate** | C4H8O3 | -H | 103.0401 | 1 | 9.36 | 4 |
| **3-D-Hydroxybutyrate** | C4H8O3 | -H | 103.0401 | 1 | 7.57 | 4 |
| **3-Hydroxyisobutyrate** | C4H8O3 | -H | 103.0401 | 1 | 7.36 | 4 |
| **3-Methyl-2-oxovaleric acid** | C6H10O3 | -H | 129.0557 | 1 | 15.58 | 4 |
| **3-Phosphoglyceric acid** | C3H7O7P | -H | 184.9857 | 1 | 13.75 | 4 |
| **4-Hydroxybutyrate** | C4H8O3 | -H | 103.0401 | 1 | 8.65 | 4 |
| **4-Hydroxyproline** | C5H9NO3 | -H | 130.051 | 1 | 1.16 | 2 |
| **5-Aminolevulinic acid** | C5H9NO3 | -H | 130.051 | 1 | 1.15 | 2 |
| **Phosphoribosyl pyrophosphate** | C5H13O14P3 | -H | 388.9445 | 1 | 16.11 | 4 |
| **Adenine** | C5H5N5 | -H | 134.0472 | 1 | 2.5 | 3 |
| **Adenosine** | C10H13N5O4 | -H | 266.0895 | 1 | 5.6 | 4 |
| **Adenosine 3-phosphate 5-phosphosulfate** | C10H15N5O13P2S | -H | 505.979 | 1 | 16.26 | 4 |
| **ADP-Ribose** | C15H23N5O14P2 | -H | 558.0644 | 1 | 13.94 | 4 |
| **cAMP** | C10H12N5O6P | -H | 328.0452 | 1 | 12.49 | 4 |
| **ADP** | C10H15N5O10P2 | -H | 426.0221 | 1 | 14.34 | 4 |
| **AICAR** | C9H15N4O8P | -H | 337.0555 | 1 | 9.86 | 4 |
| **Alanine** | C3H7NO2 | -H | 88.0404 | 1 | 1.18 | 2 |
| **alpha-Ketoglutarate** | C5H6O5 | -H | 145.0142 | 1 | 13.31 | 4 |
| **AMP** | C10H14N5O7P | -H | 346.0558 | 1 | 11.11 | 4 |
| **Arachidonic acid** | C20H32O2 | -H | 303.233 | 1 | 20.054 | 4 |
| **Ascorbic acid** | C6H8O6 | -H | 175.0248 | 1 | 14.16 | 4 |
| **Asparagine** | C4H8N2O3 | -H | 131.0462 | 1 | 1.15 | 2 |
| **Aspartate** | C4H7NO4 | -H | 132.0302 | 1 | 3.99 | 4 |
| **Asp-Asp** | C8H12N2O7 | -H | 247.0572 | 1 | 12.388 | 4 |
| **Asp-Glu** | C9H14N2O7 | -H | 261.0728 | 1 | 12.3 | 4 |
| **Asp-Gly** | C6H10N2O5 | -H | 189.0517 | 1 | 3.741 | 4 |
| **ATP** | C10H16N5O13P3 | -H | 505.9885 | 1 | 16.2 | 4 |
| **Bilirubin** | C33H36N4O6 | -H | 583.2562 | 1 | 21.71 | 4 |
| **Bisphosphoglycerate** | C3H8O10P2 | -H | 264.952 | 1 | 16.18 | 4 |

| | | | | | | |
|---|---|---|---|---|---|---|
| **CDP** | C9H15N3O11P2 | -H | 402.0109 | 1 | 13.48 | 4 |
| **CDP-ethanolamine** | C11H20N4O11P2 | -H | 445.0531 | 1 | 5.4 | 4 |
| **CDP-n-acetylneuraminic acid** | C20H31N4O16P | -H | 613.14 | 1 | 13.309 | 4 |
| **Citraconic acid** | C5H6O4 | -H | 129.0193 | 1 | 13.3 | 4 |
| **Citrate** | C6H8O7 | -H | 191.0197 | 1 | 13.92 | 4 |
| **Citrulline** | C6H13N3O3 | -H | 174.0884 | 1 | 1.18 | 2 |
| **CMP** | C9H14N3O8P | -H | 322.0446 | 1 | 8.64 | 4 |
| **Coenzyme A** | C21H36N7O16P3S | -H | 766.1079 | 1 | 16.9 | 4 |
| **Creatine** | C4H9N3O2 | -H | 130.0622 | 1 | 1.2 | 2 |
| **Creatinine** | C4H7N3O | -H | 112.0516 | 1 | 1.29 | 2 |
| **CTP** | C9H16N3O14P3 | -H | 481.9772 | 1 | 15.92 | 4 |
| **Cystathionine** | C7H14N2O4S | -H | 221.0602 | 1 | 1.15 | 2 |
| **Cytidine** | C9H13N3O5 | -H | 242.0782 | 1 | 1.62 | 2 |
| **Cytosine** | C4H5N3O | -H | 110.036 | 1 | 1.15 | 2 |
| **D-2-Hydroxyglutaric acid** | C5H8O5 | -H | 147.0299 | 1 | 12.85 | 4 |
| **dCMP** | C9H14N3O7P | -H | 306.0497 | 1 | 9.635 | 4 |
| **Dihydroorotic acid** | C5H6N2O4 | -H | 157.0255 | 1 | 6.33 | 4 |
| **Dihydroxyacetone phosphate** | C3H7O6P | -H | 168.9907 | 1 | 9.39 | 4 |
| **Dimethylglycine** | C4H9NO2 | -H | 102.0561 | 1 | 1.15 | 2 |
| **Dopamine** | C8H11NO2 | -H | 152.0717 | 1 | 1.08 | 2 |
| **dTDP** | C10H16N2O11P2 | -H | 401.0157 | 1 | 14.38 | 4 |
| **dTMP** | C10H15N2O8P | -H | 321.0493 | 1 | 10.91 | 4 |
| **dTTP** | C10H17N2O14P3 | -H | 480.982 | 1 | 16.26 | 4 |
| **FA 16:0** | C16H32O2 | -H | 255.233 | 1 | 22.34 | 3 |
| **FA 16:1** | C16H30O2 | -H | 253.2173 | 1 | 21.91 | 4 |
| **FA 18:0** | C18H36O2 | -H | 283.2643 | 1 | 23.07 | 1.5 |
| **FA 18:1** | C18H34O2 | -H | 281.2486 | 1 | 22.44 | 2 |
| **FA 18:2** | C18H32O2 | -H | 279.233 | 1 | 22.05 | 2 |
| **FA 18:3** | C18H30O2 | -H | 277.2173 | 1 | 21.77 | 4 |
| **FA 20:1** | C20H38O2 | -H | 309.2799 | 1 | 23.14 | 1 |
| **FA 20:2** | C20H36O2 | -H | 307.2643 | 1 | 22.65 | 2 |
| **FA 20:3** | C20H34O2 | -H | 305.2486 | 1 | 22.23 | 3 |
| **FA 22:2** | C22H40O2 | -H | 335.2956 | 1 | 23.34 | 1 |
| **FA 22:5** | C22H34O2 | -H | 329.2486 | 1 | 21.99 | 4 |
| **FA 22:6** | C22H32O2 | -H | 327.233 | 1 | 21.91 | 4 |
| **Farnesyl pyrophosphate** | C15H28O7P2 | -H | 381.1237 | 1 | 20.64 | 4 |
| **FAD** | C27H33N9O15P2 | -H | 784.1499 | 1 | 16.05 | 4 |
| **FMN** | C17H21N4O9P | -H | 455.0973 | 1 | 14.81 | 4 |

| | | | | | | |
|---|---|---|---|---|---|---|
| **Folic acid** | C19H19N7O6 | -H | 440.1324 | 1 | 14.23 | 4 |
| **Fructose 1,6-BP** | C6H14O12P2 | -H | 338.9888 | 1 | 14.004 | 4 |
| **Fumaric acid** | C4H4O4 | -H | 115.0037 | 1 | 13.7 | 4 |
| **GABA** | C4H9NO2 | -H | 102.0561 | 1 | 1.17 | 2 |
| **Galactonic acid** | C6H12O7 | -H | 195.051 | 1 | 4.58 | 4 |
| **gamma-Glu-Cys** | C8H14N2O5S | -H | 249.0551 | 1 | 7.486 | 4 |
| **GDP** | C10H15N5O11P2 | -H | 442.0171 | 1 | 13.83 | 4 |
| **GDP-Mannose** | C16H25N5O16P2 | -H | 604.0699 | 1 | 13.25 | 4 |
| **Gln-Glu** | C10H17N3O6 | -H | 274.1045 | 1 | 5.62 | 4 |
| **Glu-Ala** | C8H14N2O5 | -H | 217.083 | 1 | 7.534 | 4 |
| **Gluconic acid** | C6H12O7 | -H | 195.051 | 1 | 4.84 | 4 |
| **Glu-Gly** | C7H12N2O5 | -H | 203.0673 | 1 | 4.672 | 4 |
| **Glu-Ile** | C11H20N2O5 | -H | 259.1299 | 1 | 12.365 | 4 |
| **Glutaconic acid** | C5H6O4 | -H | 129.0193 | 1 | 13.69 | 4 |
| **Glutamate** | C5H9NO4 | -H | 146.0459 | 1 | 3.9 | 4 |
| **Glutamine** | C5H10N2O3 | -H | 145.0619 | 1 | 1.15 | 2 |
| **5-L-Glutamyl-taurine** | C7H14N2O6S | -H | 253.05 | 1 | 5.291 | 4 |
| **gamma-Glu-Thr** | C9H16N2O6 | -H | 247.0936 | 1 | 5.672 | 4 |
| **gamma-Glu-Val** | C10H18N2O5 | -H | 245.1143 | 1 | 9.995 | 4 |
| **Glycerol 3-phosphate** | C3H9O6P | -H | 171.0064 | 1 | 7.4 | 4 |
| **Glycine** | C2H5NO2 | -H | 74.0248 | 1 | 1.13 | 2 |
| **GMP** | C10H14N5O8P | -H | 362.0507 | 1 | 10.03 | 4 |
| **GTP** | C10H16N5O14P3 | -H | 521.9834 | 1 | 16.01 | 4 |
| **Hexose phosphate I** | C6H13O9P | -H | 259.0224 | 1 | 6.39 | 4 |
| **Hexose phosphate II** | C6H13O9P | -H | 259.0224 | 1 | 6.6 | 4 |
| **Hexose phosphate III** | C6H13O9P | -H | 259.0224 | 1 | 6.86 | 4 |
| **Hexose phosphate IV** | C6H13O9P | -H | 259.0224 | 1 | 7.06 | 4 |
| **Hexose phosphate IX** | C6H13O9P | -H | 259.0224 | 1 | 9.4 | 4 |
| **Hexose phosphate V** | C6H13O9P | -H | 259.0224 | 1 | 7.19 | 4 |
| **Hexose phosphate VI** | C6H13O9P | -H | 259.0224 | 1 | 7.41 | 4 |
| **Hexose phosphate VII** | C6H13O9P | -H | 259.0224 | 1 | 8.6 | 4 |
| **Hexose phosphate VIII** | C6H13O9P | -H | 259.0224 | 1 | 9.16 | 4 |
| **Hexose phosphate X** | C6H13O9P | -H | 259.0224 | 1 | 10.49 | 4 |
| **Histidine** | C6H9N3O2 | -H | 154.0622 | 1 | 1.04 | 2 |
| **Hydroxydecanoic acid** | C10H20O3 | -H | 187.134 | 1 | 17.32 | 4 |
| **Hydroxyhexadecanoic acid** | C16H32O3 | -H | 271.2279 | 1 | 21.58 | 4 |
| **5-Hydroxyindoleacetic acid** | C10H9NO3 | -H | 190.051 | 1 | 12.09 | 4 |
| **2-Hydroxystearic acid** | C18H36O3 | -H | 299.2592 | 1 | 22.037 | 3 |
| **Hypoxanthine** | C5H4N4O | -H | 135.0312 | 1 | 1.7 | 2 |
| **IMP** | C10H13N4O8P | -H | 347.0398 | 1 | 10.01 | 4 |

| | | | | | | |
|---|---|---|---|---|---|---|
| **Indolelactic acid** | C11H11NO3 | -H | 204.0666 | 1 | 15.53 | 4 |
| **Indoxyl sulfate** | C8H7NO4S | -H | 212.0023 | 1 | 15.5 | 4 |
| **Inosine** | C10H12N4O5 | -H | 267.0735 | 1 | 2.5 | 4 |
| **Isoleucine** | C6H13NO2 | -H | 130.0874 | 1 | 1.87 | 3 |
| **Itaconate** | C5H6O4 | -H | 129.0193 | 1 | 12.6 | 4 |
| **Ketoisovalerate** | C5H8O3 | -H | 115.0401 | 1 | 13.84 | 4 |
| **Ketoleucine** | C6H10O3 | -H | 129.0557 | 1 | 15.92 | 4 |
| **Lactate** | C3H6O3 | -H | 89.0244 | 1 | 6.918 | 4 |
| **Leucine** | C6H13NO2 | -H | 130.0874 | 1 | 2 | 3 |
| **Glutathione (reduced)** | C10H17N3O6S | -H | 306.0765 | 1 | 7.61 | 4 |
| **Glutathione (oxidized)** | C20H32N6O12S2 | -H | 611.1447 | 1 | 12.57 | 4 |
| **Malate** | C4H6O5 | -H | 133.0142 | 1 | 12.72 | 4 |
| **Malonic acid** | C3H4O4 | -H | 103.0037 | 1 | 11.72 | 4 |
| **Methionine** | C5H11NO2S | -H | 148.0438 | 1 | 1.6 | 2 |
| **2-Methylcitrate** | C7H10O7 | -H | 205.0354 | 1 | 14.38 | 4 |
| **5'-Methylthioadenosine** | C11H15N5O3S | -H | 296.0823 | 1 | 10.88 | 4 |
| **N-Acetyl-D-galactosamine 1-phosphate** | C8H16NO9P | -H | 300.049 | 1 | 7.92 | 4 |
| **N-Acetyl-1-aspartylglutamic acid** | C11H16N2O8 | -H | 303.0834 | 1 | 15.255 | 4 |
| **N-Acetyl-L-alanine** | C5H9NO3 | -H | 130.051 | 1 | 8.17 | 4 |
| **N-Acetyl-D-Glucosamine 6-Phosphate** | C8H16NO9P | -H | 300.049 | 1 | 8.198 | 4 |
| **N-Acetyl-L-glutamic acid** | C7H11NO5 | -H | 188.0564 | 1 | 13.15 | 4 |
| **N-Acetyl-L-aspartic acid** | C6H9NO5 | -H | 174.0408 | 1 | 13.05 | 4 |
| **N-Acetylglutamine** | C7H12N2O4 | -H | 187.0724 | 1 | 6.859 | 4 |
| **N-Acetyl-L-methionine** | C7H13NO3S | -H | 190.0543 | 1 | 13.1 | 4 |
| **N-Acetylneuraminic acid** | C11H19NO9 | -H | 308.0987 | 1 | 5.717 | 4 |
| **N-Acetyl-L-phenylalanine** | C11H13NO3 | -H | 206.0823 | 1 | 15.61 | 4 |
| **N-Acetylserine** | C5H9NO4 | -H | 146.0459 | 1 | 5.9 | 4 |
| **N-Acetyltryptophan** | C13H14N2O3 | -H | 245.0932 | 1 | 15.56 | 4 |
| **NAD+** | C21H27N7O14P2 | -H | 662.1018 | 1 | 8.61 | 4 |
| **NADH** | C21H29N7O14P2 | -H | 664.1175 | 1 | 14.41 | 4 |
| **NADP+** | C21H28N7O17P3 | -H | 742.0682 | 1 | 13.81 | 4 |
| **NADPH** | C21H30N7O17P3 | -H | 744.0838 | 1 | 16.18 | 4 |
| **Carglumic acid** | C6H10N2O5 | -H | 189.0517 | 1 | 12.61 | 4 |

| | | | | | | |
|---|---|---|---|---|---|---|
| **N-Glycolylneuraminic acid** | C11H19NO10 | -H | 324.0936 | 1 | 4.95 | 4 |
| **O-Acetylserine** | C5H9NO4 | -H | 146.0459 | 1 | 6.37 | 4 |
| **O-Phosphoethanolamine** | C2H8NO4P | -H | 140.0118 | 1 | 1.65 | 3 |
| **Ophthalmate** | C11H19N3O6 | -H | 288.1201 | 1 | 7.85 | 4 |
| **Orotic acid** | C5H4N2O4 | -H | 155.0098 | 1 | 8.15 | 4 |
| **Orotidine** | C10H12N2O8 | -H | 287.0521 | 1 | 7.221 | 4 |
| **Pantothenic acid** | C9H17NO5 | -H | 218.1034 | 1 | 11.17 | 4 |
| **Pentose phosphate I** | C5H11O8P | -H | 229.0119 | 1 | 6.82 | 4 |
| **Pentose phosphate III** | C5H11O8P | -H | 229.0119 | 1 | 8 | 4 |
| **Pentose phosphate IV** | C5H11O8P | -H | 229.0119 | 1 | 8.7 | 4 |
| **Pentose phosphate V** | C5H11O8P | -H | 229.0119 | 1 | 10.33 | 4 |
| **Pentose phosphate II** | C5H11O8P | -H | 229.0119 | 1 | 7.8 | 4 |
| **Phenylalanine** | C9H11NO2 | -H | 164.0717 | 1 | 3.75 | 4 |
| **Phenyllactic acid** | C9H10O3 | -H | 165.0557 | 1 | 15.61 | 4 |
| **Phosphocreatine** | C4H10N3O5P | -H | 210.0285 | 1 | 12.52 | 4 |
| **Phosphoenolpyruvate** | C3H5O6P | -H | 166.9751 | 1 | 14.29 | 4 |
| **Phosphoserine** | C3H8NO6P | -H | 184.0016 | 1 | 8.79 | 4 |
| **Hydroxyphenyllactic acid** | C9H10O4 | -H | 181.0506 | 1 | 12.06 | 4 |
| **Proline** | C5H9NO2 | -H | 114.0561 | 1 | 1.22 | 2 |
| **Pyridoxal** | C8H9NO3 | -H | 166.051 | 1 | 2.06 | 3 |
| **Pyridoxal 5'-phosphate** | C8H10NO6P | -H | 246.0173 | 1 | 13.95 | 4 |
| **Pyridoxine** | C8H11NO3 | -H | 168.0666 | 1 | 2.13 | 3 |
| **Pyroglutamic acid** | C5H7NO3 | -H | 128.0353 | 1 | 7.065 | 4 |
| **Pyruvate** | C3H4O3 | -H | 87.0088 | 1 | 8.64 | 4 |
| **Riboflavin** | C17H20N4O6 | -H | 375.131 | 1 | 11.9 | 4 |
| **SAH** | C14H20N6O5S | -H | 383.1143 | 1 | 4.55 | 4 |
| **Sedoheptulose 7-phosphate** | C7H15O10P | -H | 289.033 | 1 | 7.045 | 4 |
| **Serine** | C3H7NO3 | -H | 104.0353 | 1 | 1.14 | 2 |
| **S-Lactoylglutathione** | C13H21N3O8S | -H | 378.0977 | 1 | 13.616 | 4 |
| **Succinate** | C4H6O4 | -H | 117.0193 | 1 | 12.05 | 4 |
| **Taurine** | C2H7NO3S | -H | 124.0074 | 1 | 1.18 | 2 |
| **Threonic acid** | C4H8O5 | -H | 135.0299 | 1 | 5.01 | 4 |
| **Threonine** | C4H9NO3 | -H | 118.051 | 1 | 1.17 | 2 |
| **Thymidine** | C10H14N2O5 | -H | 241.083 | 1 | 4.4 | 4 |
| **Tryptophan** | C11H12N2O2 | -H | 203.0826 | 1 | 7.422 | 4 |
| **Tyrosine** | C9H11NO3 | -H | 180.0666 | 1 | 1.95 | 3 |
| **UDP** | C9H14N2O12P2 | -H | 402.9949 | 1 | 13.82 | 4 |

| | | | | | | |
|---|---|---|---|---|---|---|
| **UDP-alpha-D-glucuronic acid** | C15H22N2O18P2 | -H | 579.027 | 1 | 15.88 | 4 |
| **UDP-hexose** | C15H25N2O17P2 | -H | 566.0556 | 1 | 13.29 | 4 |
| **UDP-N-acetylhexosamine** | C17H27N3O17P2 | -H | 606.0743 | 1 | 13.37 | 4 |
| **UMP** | C9H13N2O9P | -H | 323.0286 | 1 | 9.65 | 4 |
| **Uracil** | C4H4N2O2 | -H | 111.02 | 1 | 1.49 | 2 |
| **Ureidosuccinic acid** | C5H8N2O5 | -H | 175.036 | 1 | 12.58 | 4 |
| **Uric acid** | C5H4N4O3 | -H | 167.0211 | 1 | 5.07 | 4 |
| **Uridine** | C9H12N2O6 | -H | 243.0623 | 1 | 1.95 | 3 |
| **UTP** | C9H15N2O15P3 | -H | 482.9613 | 1 | 16.09 | 4 |
| **Valine** | C5H11NO2 | -H | 116.0717 | 1 | 1.35 | 2 |
| **Xanthine** | C5H4N4O2 | -H | 151.0261 | 1 | 1.7 | 2 |
| **Xanthosine** | C10H12N4O6 | -H | 283.0684 | 1 | 7.28 | 4 |

### Uncropped Western blots

#### From Fig. 4g

Total LRRK2

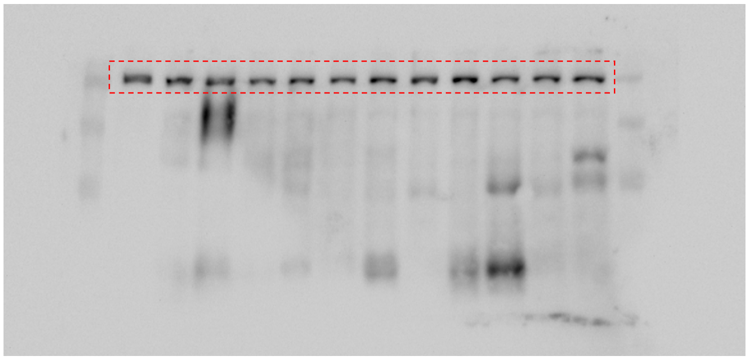

pSer935-LRRK2

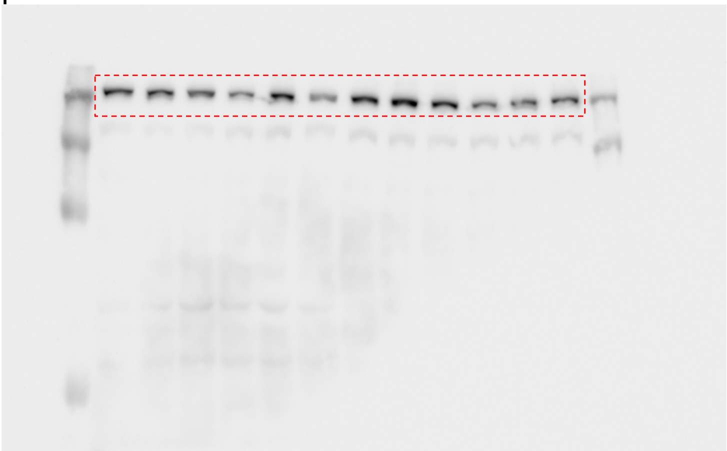

Total Rab12

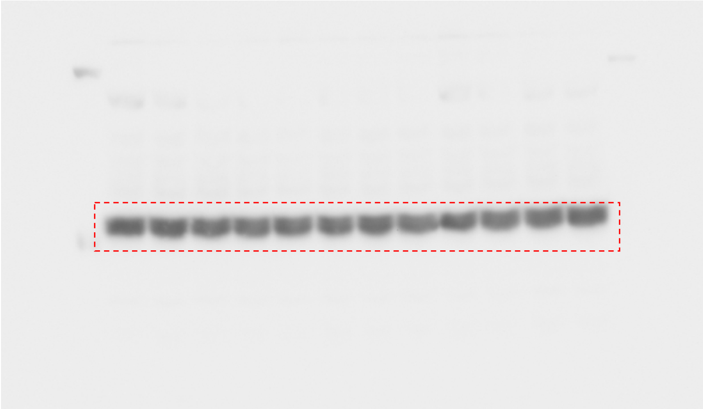

pSer106-Rab12

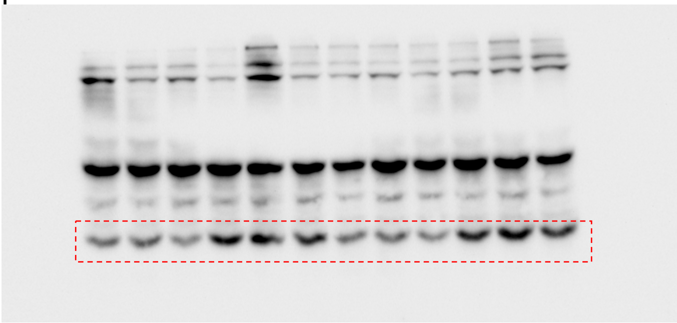

Actin

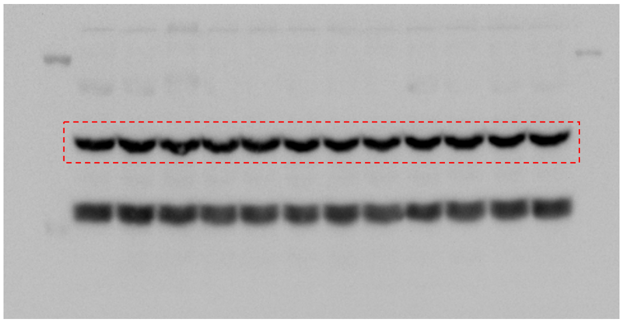

**From Fig. 4h**

Total LRRK2

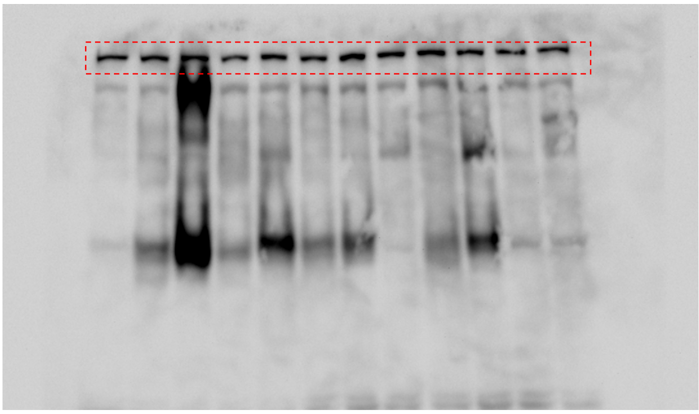

pSer935-LRRK2

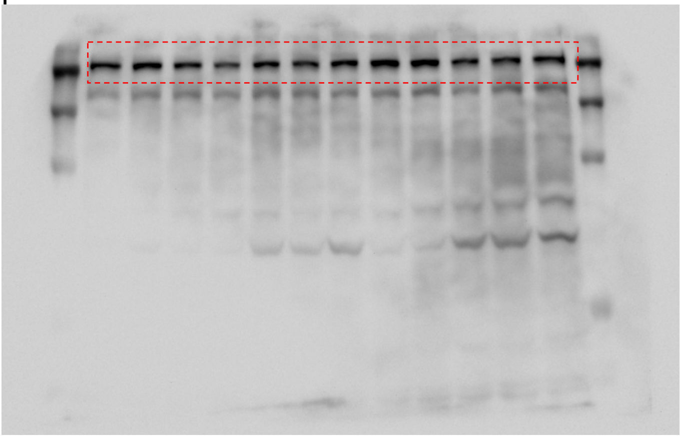

Total Rab12

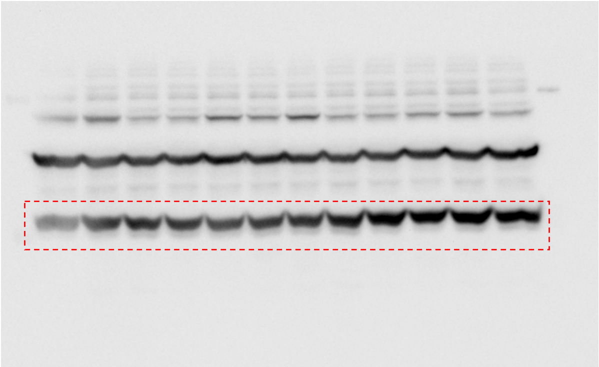

pSer106-Rab12

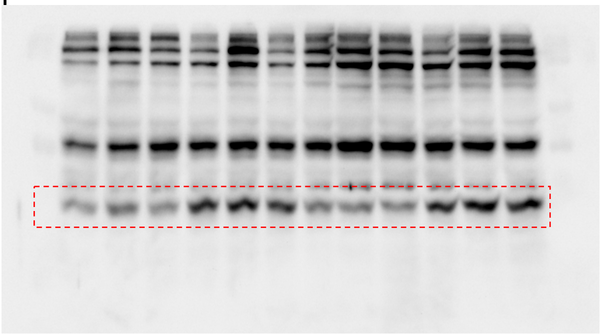

Actin

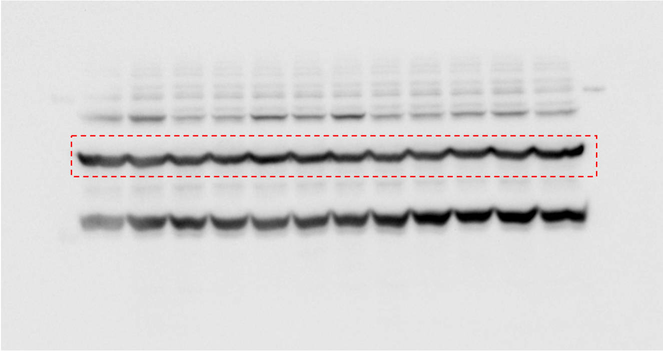

**From Fig. 9b**

Thymidine Phosphorylase (TP)

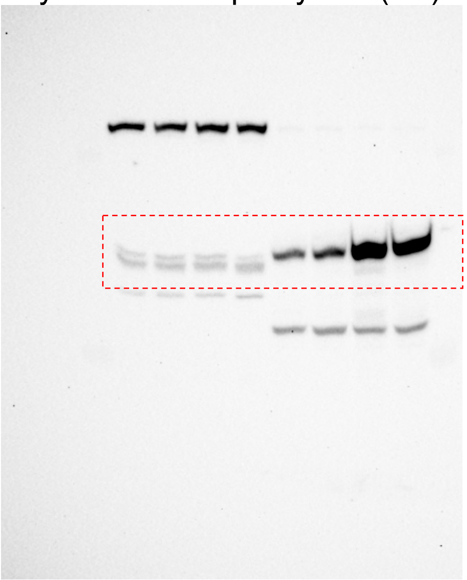

Actin

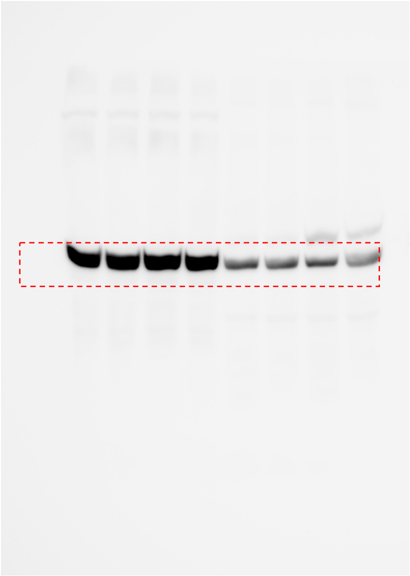

**From Fig. 9f**

TP (upper band) / Actin (lower band)

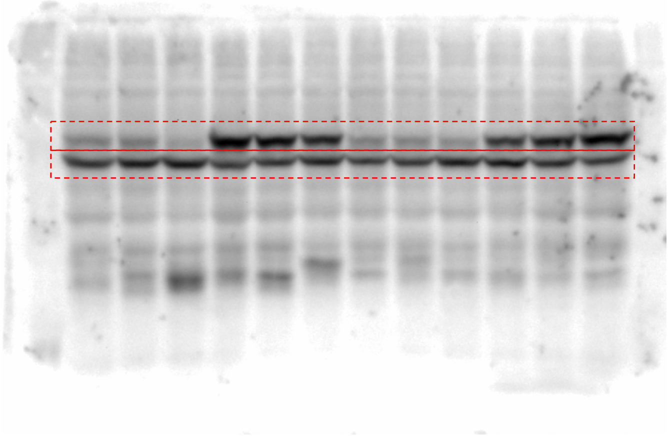

Total LRRK2

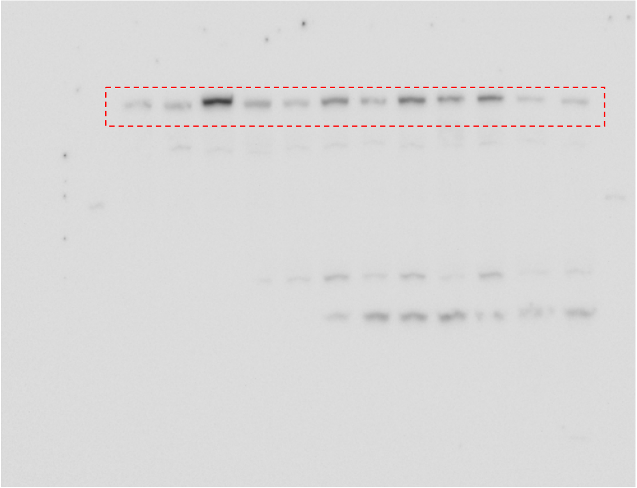

pSer1292-LRRK2

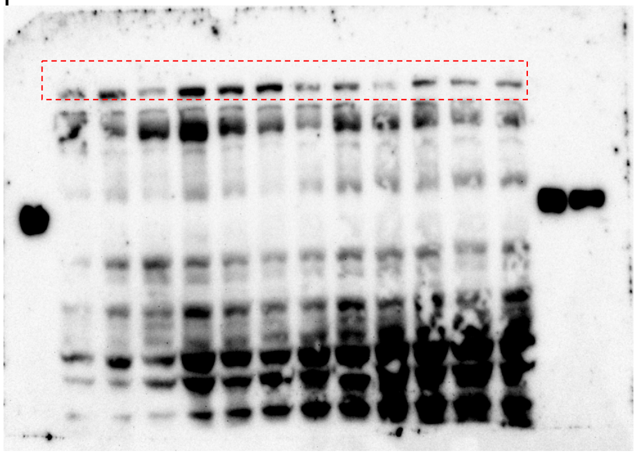

pSer935-LRRK2

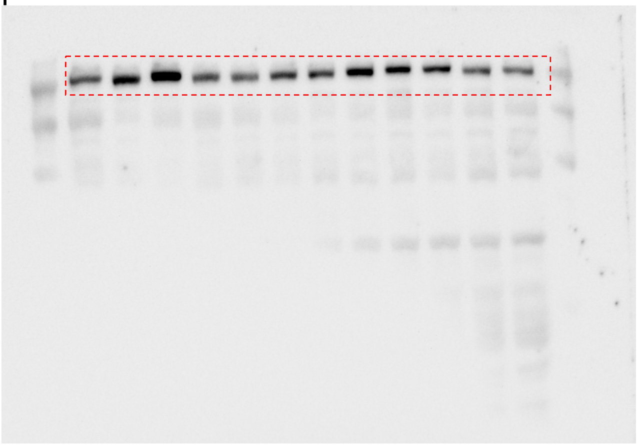

Total Rab10

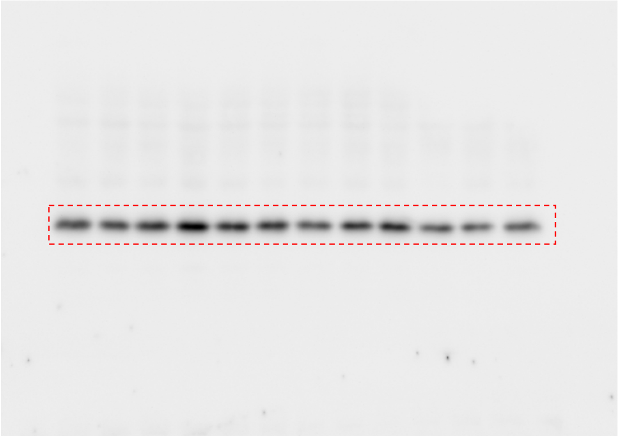

pThr73-Rab10

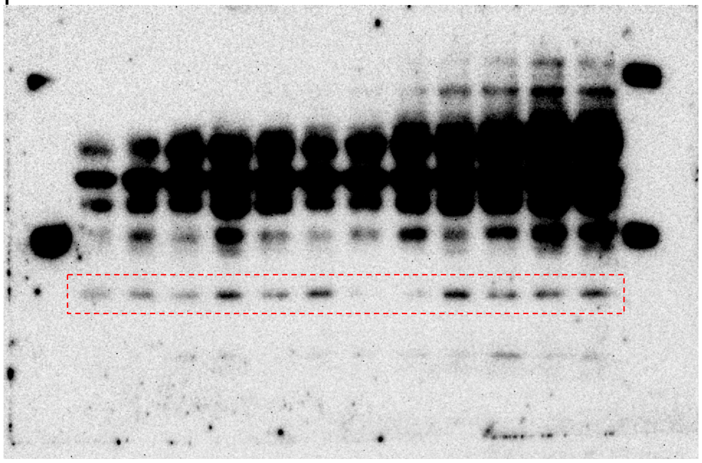

Total Rab12

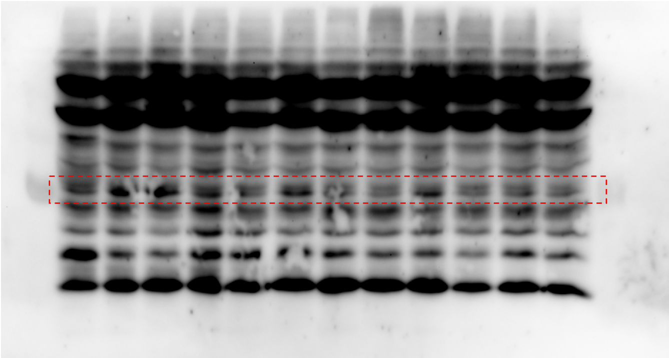

pSer106-Rab12

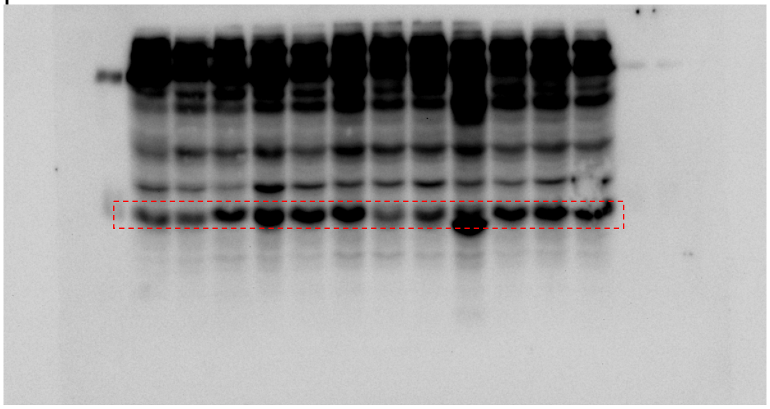

**From Fig. 10x**

Total LRRK2

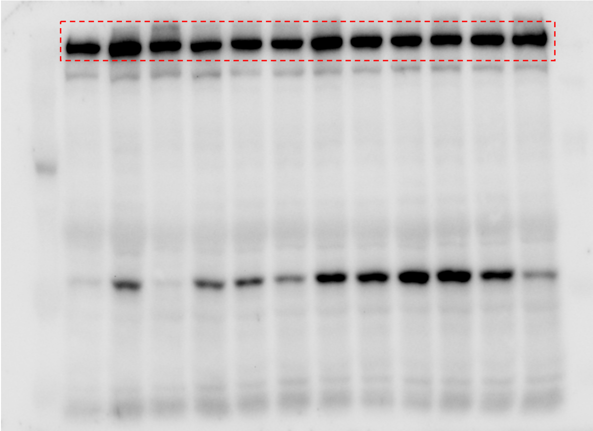

pSer935-LRRK2

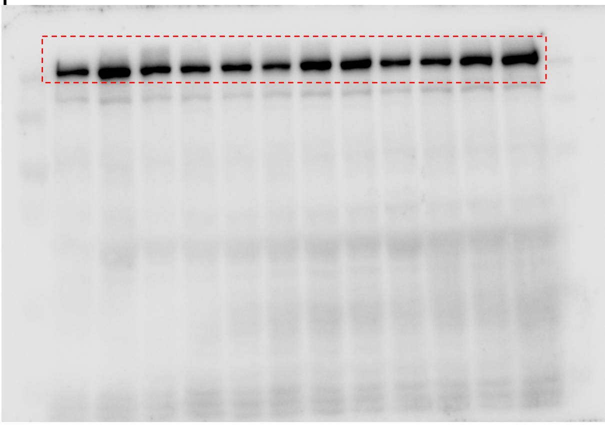

pSer1292-LRRK2

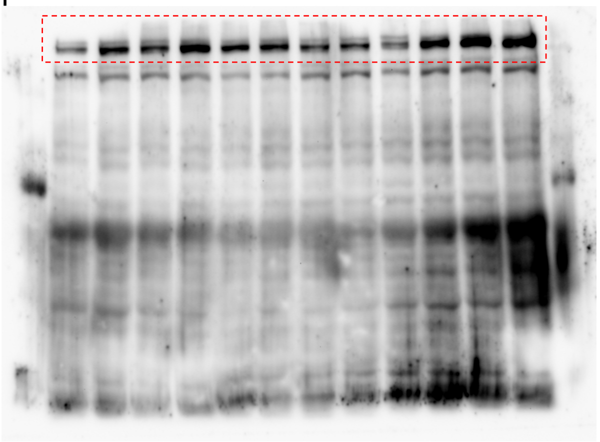

Actin

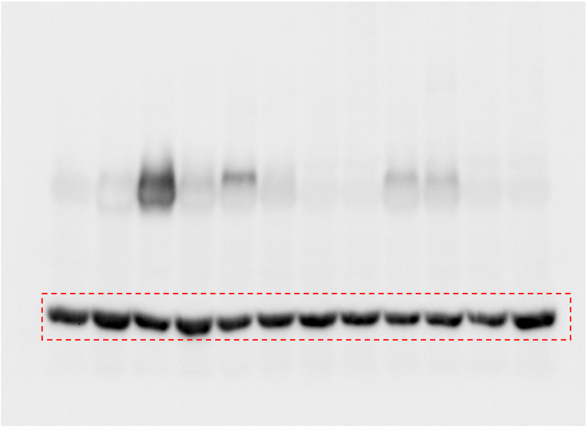

Total Rab8a

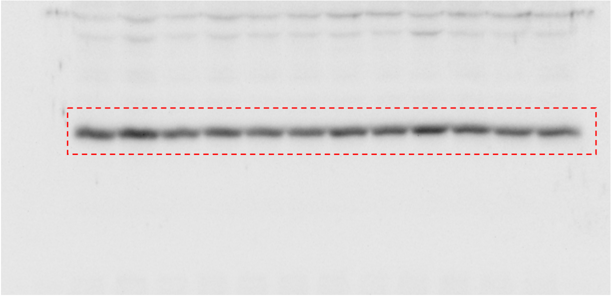

pThr72-Rab8a

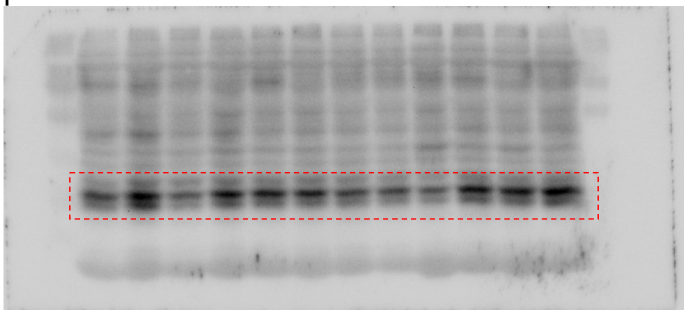

Total Rab10

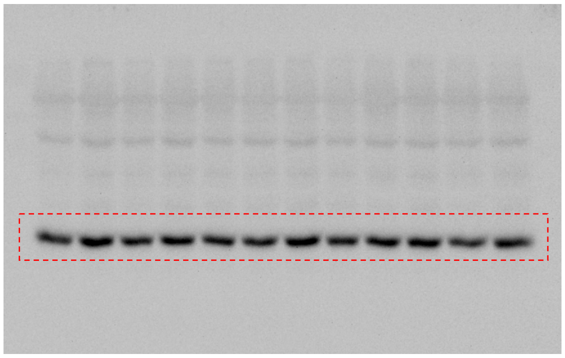

pThr73-Rab10

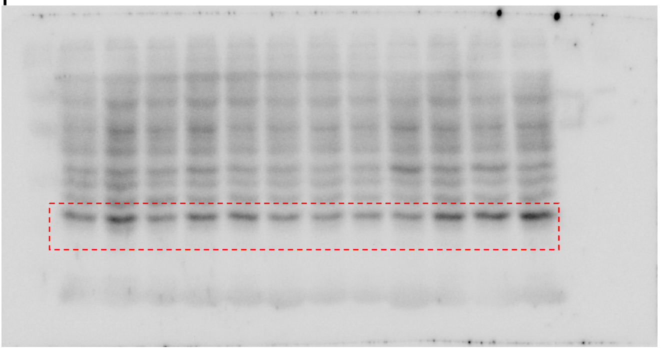

Total Rab12

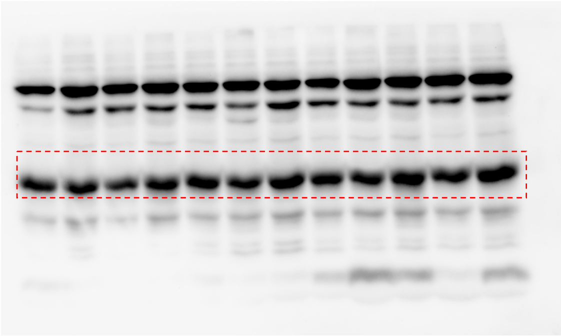

pSer106-Rab12

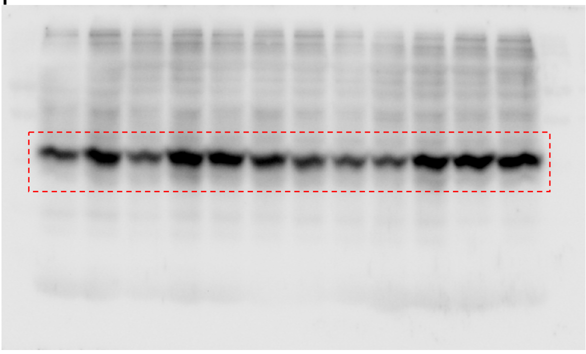

Actin

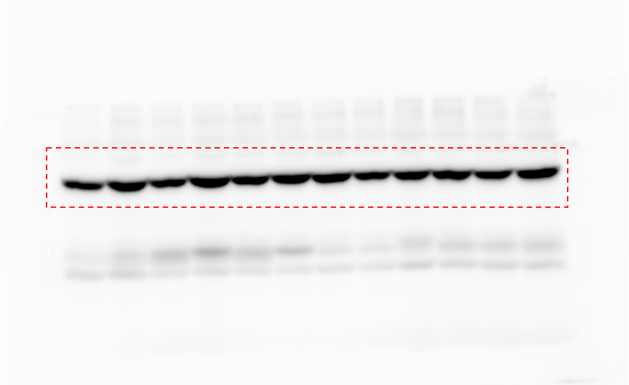
